# How can graph theory inform the dual-stream model of speech processing? a resting-state fMRI study of post-stroke aphasia

**DOI:** 10.1101/2023.04.17.537216

**Authors:** Haoze Zhu, Megan C. Fitzhugh, Lynsey M. Keator, Lisa Johnson, Chris Rorden, Leonardo Bonilha, Julius Fridriksson, Corianne Rogalsky

## Abstract

The dual-stream model of speech processing has been proposed to represent the cortical networks involved in speech comprehension and production. Although it is arguably the prominent neuroanatomical model of speech processing, it is not yet known if the dual-stream model represents actual intrinsic functional brain networks. Furthermore, it is unclear how disruptions after a stroke to the functional connectivity of the dual-stream model’s regions are related to specific types of speech production and comprehension impairments seen in aphasia. To address these questions, in the present study, we examined two independent resting-state fMRI datasets: (1) 28 neurotypical matched controls and (2) 28 chronic left-hemisphere stroke survivors with aphasia collected at another site. Structural MRI, as well as language and cognitive behavioral assessments, were collected. Using standard functional connectivity measures, we successfully identified an intrinsic resting-state network amongst the dual-stream model’s regions in the control group. We then used both standard functional connectivity analyses and graph theory approaches to determine how the functional connectivity of the dual-stream network differs in individuals with post-stroke aphasia, and how this connectivity may predict performance on clinical aphasia assessments. Our findings provide strong evidence that the dual-stream model is an intrinsic network as measured via resting-state MRI, and that weaker functional connectivity of the hub nodes of the dual-stream network defined by graph theory methods, but not overall average network connectivity, is weaker in the stroke group than in the control participants. Also, the functional connectivity of the hub nodes predicted specific types of impairments on clinical assessments. In particular, the relative strength of connectivity of the right hemisphere’s homologues of the left dorsal stream hubs to the left dorsal hubs versus right ventral stream hubs is a particularly strong predictor of post-stroke aphasia severity and symptomology.

## Introduction

In the past two decades, the dual-stream model of speech processing (Hickok & Poeppel, 2000, 2004, 2007) has, arguably, emerged as the prominent functional anatomical model of speech perception and production (Crinion & Price, 2005; Humphries, Willard, Buchsbaum, & Hickok, 2001; Keator et al., 2022; Rogalsky & Hickok, 2009; Spitsyna, Warren, Scott, Turkheimer, & Wise, 2006; Vandenberghe, Nobre, & Price, 2002). During this same time, the connectivity of brain regions with similar response properties has emerged as a valuable predictor of behavioral performance and neurological disease (Battistella et al., 2020; Ferguson et al., 2019; Fitzhugh, Hemesath, Keator et al., 2021; Schaefer, Baxter, & Rogalsky, 2019; Walsh, Baxter, Smith, & Braden, 2019; Zhang et al., 2021; Zhu et al., 2016). However, it remains unclear if the dual-stream model of speech processing represents an intrinsic, coherent brain network, and, if so, the nature of its organizational structure and how damage to one part of the network affects overall network function and behavioral speech performance remains unclear. Resting-state fMRI is a particularly promising, emerging tool to explore the dual-stream network as it relates to post-stroke aphasia severity and specific impairments.

It is well established that low-frequency BOLD signal fluctuations at rest indicate coherent language-related neural activity (Battistella et al., 2020; Binder et al., 1999; Tie et al., 2014; Xu, Huang, Cui, & Yu, 2020; Zhao, Ralph, & Halai, 2018). For instance, a recent study by Battistella and colleagues (2020) identified several language-related networks with resting-state fMRI in young neurotypical adults, including a dorsal articulatory-phonological and ventral semantic network. A seed-voxel based method used some critical nodes in the language-related task such as inferior frontal gyrus from an activation peak of fluency task (Heim, Eickhoff, & Amunts, 2008). In addition, another two functional connectivity studies have also found significant functional connectivity across language-related areas in control subjects, such as sensorimotor areas and expressive and receptive language regions. (Cordes et al., 2000; Hampson, Peterson, Skudlarski, Gatenby, & Gore, 2002). Based on the findings above, there is strong evidence that nodes of language networks are connected at rest, however the architecture of these networks at rest have not been well-mapped, particularly in older adult control subjects and stroke survivors.

The studies mentioned above typically represented a network with the voxel or ROI- based functional connectivity which indicated the higher correlation regions with the critical nodes or the correlations between some important cortical areas. In addition to the traditional functional connectivity to characterize networks in resting-state fMRI, graph-theoretical approaches may be a powerful method to better understand the network properties of the dual-stream model’s regions because it can identify the hub region which play a pivotal role in the network organization and some network level properties which could not be reflected using the simple functional connectivity (He, Chen, & Evans, 2008; He & Evans, 2010). Typically, graph-theoretical analyses consist of a series of nodes and edges corresponding to the constituent units and interaction of a network (Barabási, 2012; Boccaletti, Latora, Moreno, Chavez, & Hwang, 2006; Liao, Vasilakos, & He, 2017). Unlike the traditional functional connectivity focusing on the relationship between a few regions, graph theory tends to describe the organization of the whole network or the local node cluster. After building a graph, hub regions can be defined as the average shortest path length. The shortest path length is the number of edges comprising the shortest path between any pair of nodes; the average shortest path length *L_i_* is the mean of n-1 minimum pathway between the index node and all other nodes in the networks (Achard, Salvador, Whitcher, Suckling, & Bullmore, 2006).

A few studies related to language in adult control subject and children have applied graph theory to depict the interactions between brain regions known to be involved in speech processing, with greater performance on various language measures being related not just to increased functional connectivity, but to increased connectivity of hub regions, as defined by graph theory approaches (Muller & Meyer, 2014; Xiao, Friederici, Margulies, & Brauer, 2016). In the area of degenerative brain disease, hub regions have been shown to be more vulnerable (Achard et al., 2006; Albert, Jeong, & Barabási, 2000; Bassett & Bullmore, 2006), in that damage to or disruptions of hub regions are typically more correlated with more severe neurocognitive impairments than damage or disruption to non-hub regions of a network (He et al., 2008; Roger et al., 2019; Stam et al., 2009). However, to our knowledge no previous studies have examined the functional network connectivity of the dual-stream model using graph theory, and only a few studies have investigated stroke-induced language impairments using graph theory (Bohland, Kapse, & Kiran, 2014; Duncan & Small, 2016; Mazrooyisebdani, A. Nair, Garcia-ramos, & Prabhakaran, 2018). These previous studies found some network level properties were changed and related to the deficiency or recovery of post-stroke aphasia. However, the discussion of functional connectivity between the hubs was absent. By moving beyond the more typical functional connectivity analyses and using graph theory, we will be able to examine network organization, and reorganization in stroke survivors, in new ways that can potentially help us better understand post-stroke compensation and language impairments.

In the present study, we not only extend the study of the dual-stream model to resting-state fMRI, but also implement graph theory approaches in both stroke survivors and neurotypical control subjects to characterize how the dual-stream network differs due to stroke and is related to language abilities post-stroke. We have three hypotheses: 1) The neurotypical control participants will exhibit strong resting-state functional connectivity patterns across dual-stream nodes that have been defined by previous task-based findings; 2) The stroke survivors, in comparison to the control group, will exhibit functional connectivity and graph theory parameter differences, and 3) In stroke survivors, differences to the hub regions defined by graph theory will predict impairments in our behavioral language measures.

## Methods

### 2.1 Participants

#### Stroke Group

29 chronic left hemisphere stroke survivors were recruited and tested at the University of South Carolina. One of these stroke participants’ data was removed from the present study’s analyses because of substantial head motion during scanning. Remaining stroke survivors range in age from 35 to 78 years (*M*=60.00, *sd*=11.47). Inclusion criteria included chronic stroke (<6 months prior to testing), right-handed pre-stroke, native speakers of American English, 18+ years of age, and with no history of neurological disease, head trauma, or psychiatric disturbances prior to their stroke (self-report and supported by review of MRI scans by a clinical neurological radiologist). Aphasia classification was determined by the Western Aphasia Battery (Kertesz, 2007); each stroke participant’s aphasia diagnosis is reported in Table 1. Also, the lesion overlap map concludes the lesion location for the stroke group.

**Table 1.**
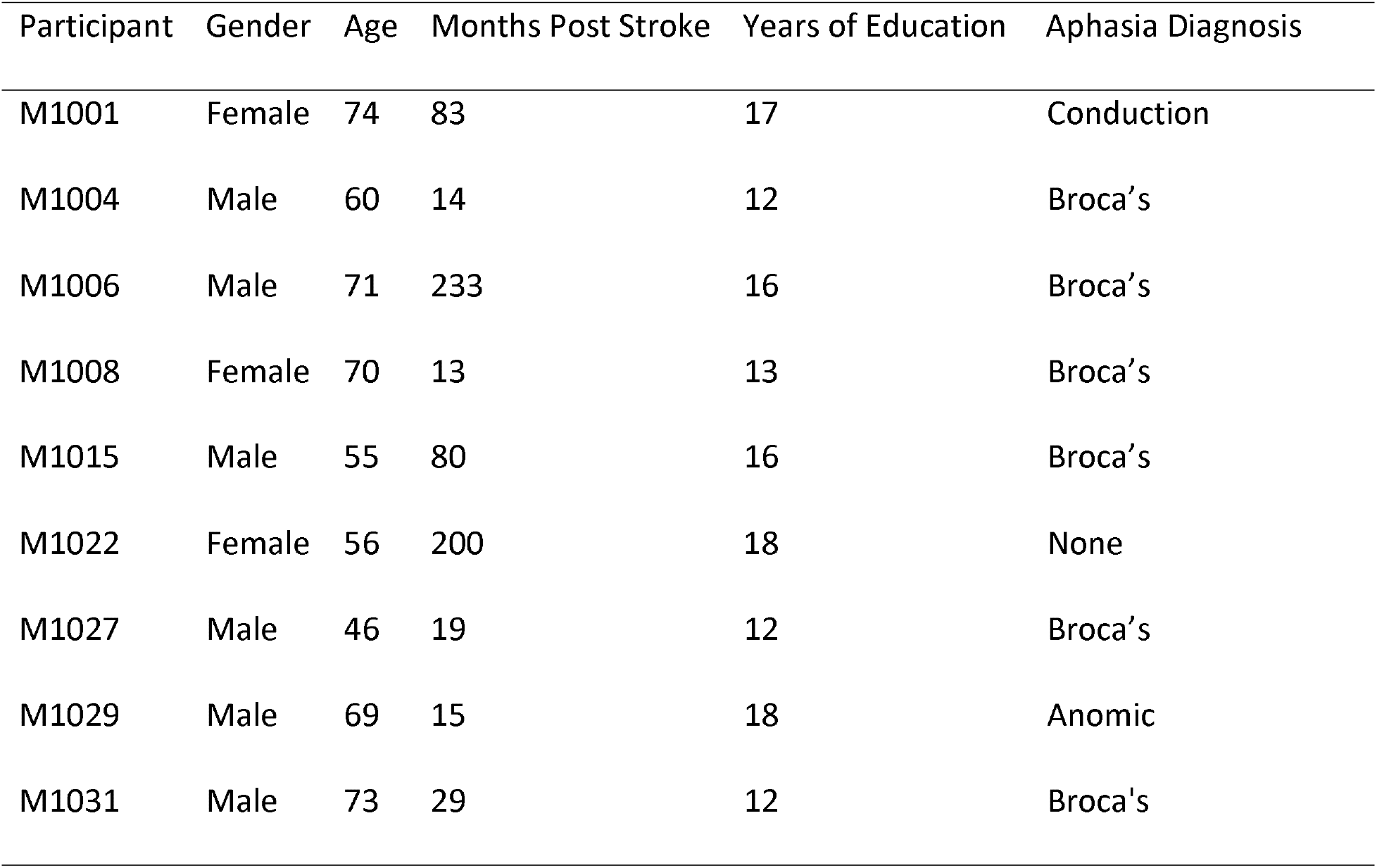
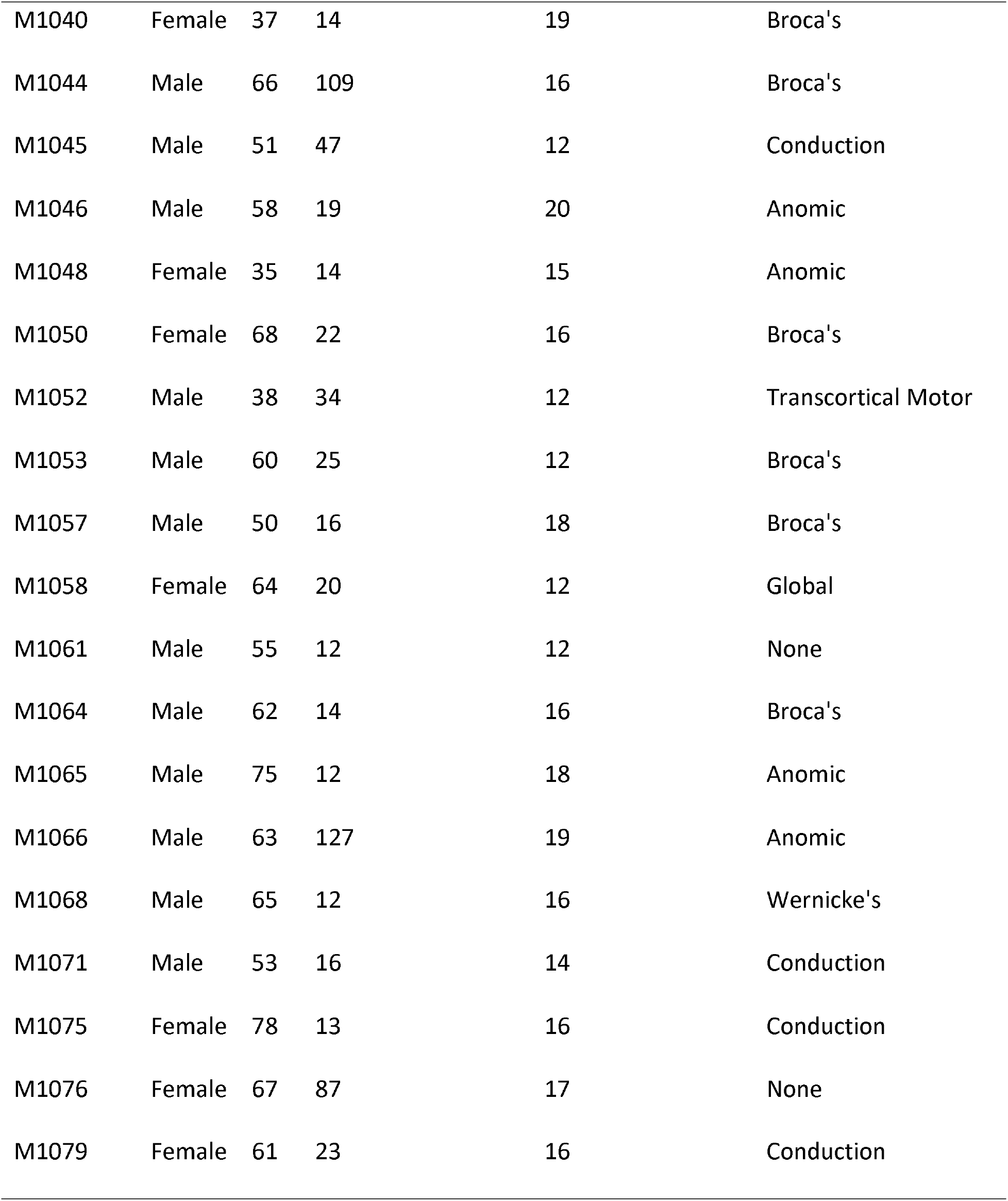
Demographics of stroke group

| Participant | Gender | Age | Months Post Stroke | Years of Education | Aphasia Diagnosis |
| --- | --- | --- | --- | --- | --- |
| M1001 | Female | 74 | 83 | 17 | Conduction |
| M1004 | Male | 60 | 14 | 12 | Broca's |
| M1006 | Male | 71 | 233 | 16 | Broca's |
| M1008 | Female | 70 | 13 | 13 | Broca's |
| M1015 | Male | 55 | 80 | 16 | Broca's |
| M1022 | Female | 56 | 200 | 18 | None |
| M1027 | Male | 46 | 19 | 12 | Broca's |
| M1029 | Male | 69 | 15 | 18 | Anomic |
| M1031 | Male | 73 | 29 | 12 | Broca's |
| M1040 | Female | 37 | 14 | 19 | Broca's |
| M1044 | Male | 66 | 109 | 16 | Broca's |
| M1045 | Male | 51 | 47 | 12 | Conduction |
| M1046 | Male | 58 | 19 | 20 | Anomic |
| M1048 | Female | 35 | 14 | 15 | Anomic |
| M1050 | Female | 68 | 22 | 16 | Broca's |
| M1052 | Male | 38 | 34 | 12 | Transcortical Motor |
| M1053 | Male | 60 | 25 | 12 | Broca's |
| M1057 | Male | 50 | 16 | 18 | Broca's |
| M1058 | Female | 64 | 20 | 12 | Global |
| M1061 | Male | 55 | 12 | 12 | None |
| M1064 | Male | 62 | 14 | 16 | Broca's |
| M1065 | Male | 75 | 12 | 18 | Anomic |
| M1066 | Male | 63 | 127 | 19 | Anomic |
| M1068 | Male | 65 | 12 | 16 | Wernicke's |
| M1071 | Male | 53 | 16 | 14 | Conduction |
| M1075 | Female | 78 | 13 | 16 | Conduction |
| M1076 | Female | 67 | 87 | 17 | None |
| M1079 | Female | 61 | 23 | 16 | Conduction |

#### Neurotypical control group

Twenty-eight neurotypical adults ranging in age from 20 to 79 years (*M*=58.79, *sd*=18.72) who were also right-handed, native speakers of American English, 18+ years of age, a minimum score of 27 on the Mini-Mental State Exam (Tombaugh & McIntyre, 1992), with no history of neurological disease, head trauma, or psychiatric disturbances were recruited from the greater Phoenix, Arizona area. How our stroke and control groups match in relation to age, gender, and education are depicted in Table 2.

**Table 2.** Comparison of demographics of stroke and control groups.

|  | Stroke(n=28) | Control(n=28) | Group Comparison Statistic |
| --- | --- | --- | --- |
| Age: mean years (sd) | 60.00(11.47) | 58.79(18.72) | $t(54) = 0.29, p = 0.77$ |
| Gender: male/female | 18/10 | 14/14 | $\chi^2(1) = 1.1667, p = 0.28$ |
| Education: mean years (sd) | 15.36(2.60) | 17.18(3.01) | $t(54) = -2.42, p = 0.02$ |

### 2.2 Image acquisition

#### Control group

MRI data of the control group was acquired on a 3T Phillips Ingenia MRI scanner equipped with a 32 channel radiofrequency head coil located at the Keller Center for Imaging Innovation at the Barrow Neurological Institute in Phoenix, Arizona. A T1 image was collected with the following parameters: FOV = 270 × 252, TR = 6.74 s, TE = 3.10 ms, flip angle = 9, voxel size = 1 × 1 × 1 mm. Resting-state fMRI data were collected using single-shot EPI with following parameters: one 10-min run, 197 total volumes, TR = 3000 ms, FOV = 217 × 217, matrix = 64 × 62, 3.39 mm slice thickness, in-plane resolution = 3.39 × 3.39 mm.

#### Stroke group

MRI data of the stroke group was acquired on a 3T Siemens scanner at Prisma Health Richland Hospital. A T2 structural image was collected with the following parameters: TE = 57 ms, image size: [176 256 256], voxel size = 1 × 1 × 1 mm. Resting-state fMRI data were collected using EPI with following parameters: one 11-min run, 427 total volumes, TR = 1650 ms, Percent Phase FOV = 100, 2 mm slice thickness. Although the stroke and control groups’ data were acquired from different scanners, there is evidence that inter-scanner variability effects on functional connectivity, such as we use here, is limited (Noble et al., 2017; Marek et al., 2019; Belleau et al., 2020).

### 2.3 Behavioral data methods

The language abilities of the stroke group were evaluated by administration of the Western Aphasia Battery (WAB) (Kertesz, 2007). The WAB’s aphasia quotient indicating overall aphasia severity, as well as the scores of the following subtests were used in subsequent analyses: Spontaneous Speech, Auditory Verbal Comprehension, Repetition, and Naming and Word Finding.

### 2.4 Data analysis

#### 2.4.1 MRI data Preprocessing

All resting-state fMRI data were preprocessed using Statistical Parametric Mapping 12 (https://www.fil.ion.ucl.ac.uk/spm). The first two time points of each run were discarded to ensure that the magnetization reached a steady state and the subjects adapted to the environment. Slice timing adjusted to compensate the interleaved acquisition in the remaining 187 volumes for control and 427 volumes for stroke survivors. Next, realignment was conducted to correct head motion using the six standard head motion parameters. Then the structural image (i.e., T1-weighted image for the control group and T2-weighted image for the stroke group) was reoriented to the mean functional image. Diffeomorphic anatomical registration through exponentiated Lie algebra normalization (DARTEL) (Ashburner 2007) was used to segment the structural image to white matter, grey matter, cerebral spinal fluid and normalize it to Montreal Neurological Institute (MNI) template. Using the normal parameters of the structural image generated by DARTEL, the functional images were spatially normalized to MNI space. Also, nuisance covariates including WM signal, CSF signal, and head motion parameters were regressed out (Fox 2005, 2006) from the functional signal. We then applied band-pass filtering between 0.01 and 0.1Hz and spatial smoothing with an 8mm FWHM Gaussian kernel to facilitate group analyses. For the stroke group’s fMRI data processing, we also added a cost function masking step (Andersen, Rapcsak, & Beeson, 2010; Brett, Leff, Rorden, & Ashburner, 2001) to avoid the risk that stretched or altered peri-lesion tissue would impair normalization. The mask used in this step was generated from a manual demarcation of the chronic stroke lesion as seen on the T2-weighted images by researchers trained in neuroanatomy and lesion mapping procedures.

#### 2.4.2 Identification of nodes in the dual-stream network

There are several meta-analysis studies to delineate left dominant regions that involves in the languages processing including left frontal, temporal, and parietal lobes (Dronkers, 2011; Margulies & Petrides, 2013; Mesulam et al., 2014; Price, 2012; Vigneau et al., 2006). These studies clearly showed the critical language-related regions such as Broca’s area, motor area, Wernicke’s area, Sylvian fissure which is in line with the dual stream model. To depict the comprehensive language processing area, we used 32 nodes from previous task-based fMRI study in the left hemisphere which used three contrasts including sentence production, listening and reading with corresponding word-list reference tasks (Labache et al., 2019). These regions are highly consistent with typical language areas. Out of the LaBache coordinates, precentral sulcus, superior frontal gyrus, inferior frontal sulcus, two Broca’s areas nodes and one anterior insula node were classified as dorsal, superior temporal gyrus, two middle temporal gyrus nodes, four superior temporal sulcus nodes, supramarginal gyrus and angular gyrus were classified as ventral based on location. All the nodes in the left hemisphere are listed in Table 3.

**Table 3.**
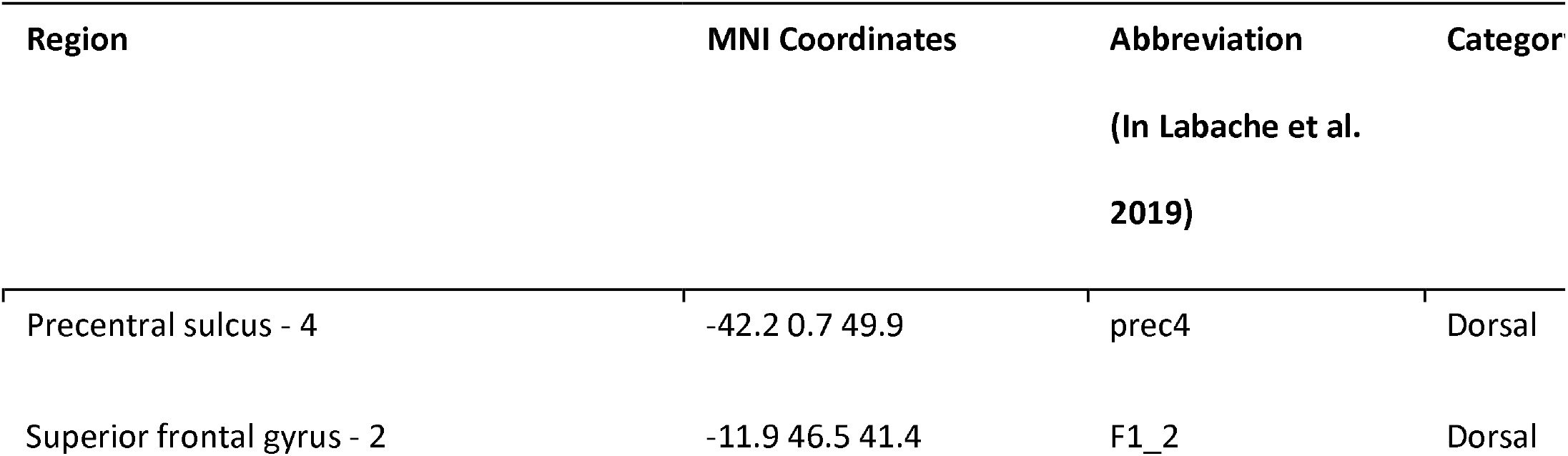
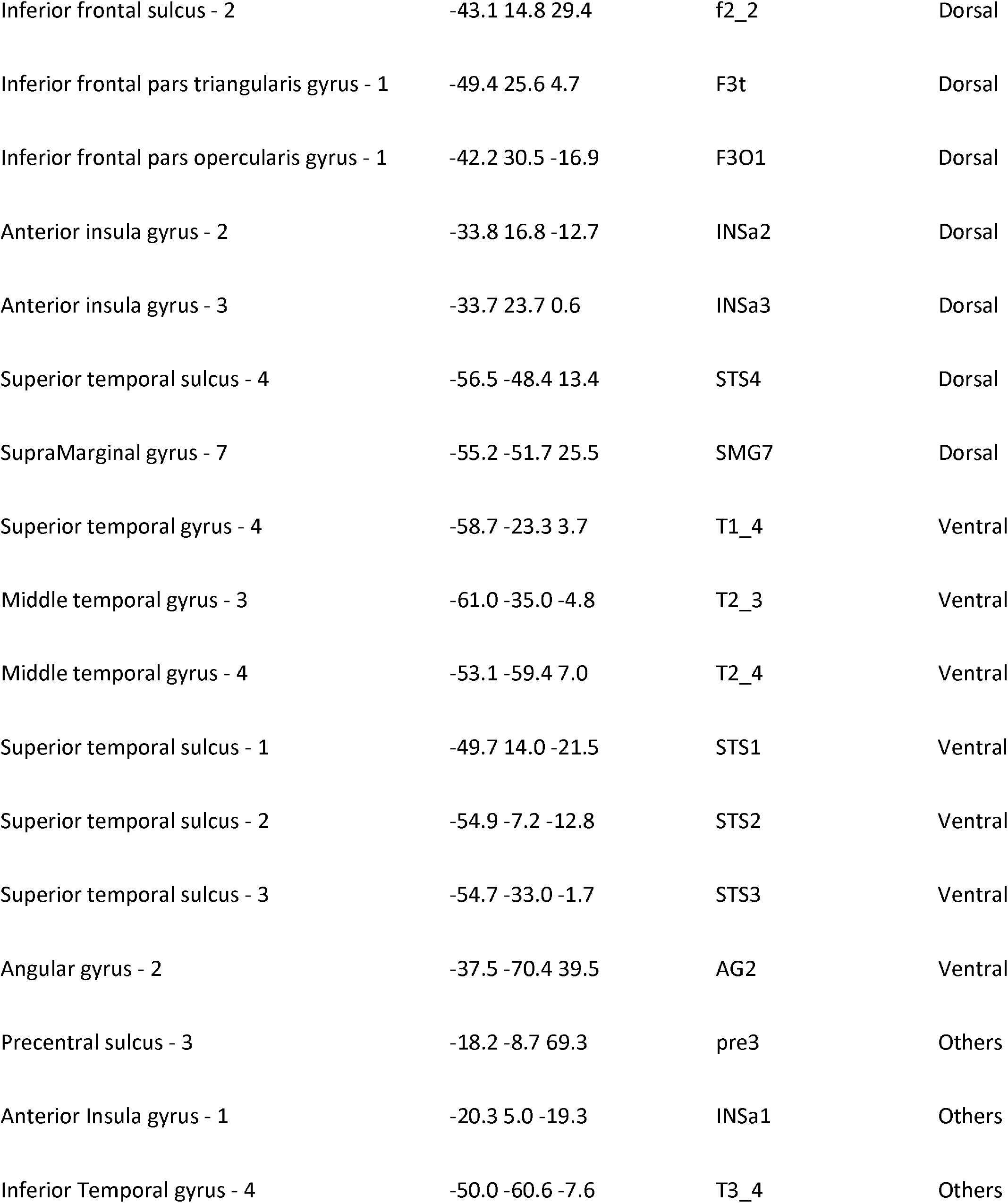
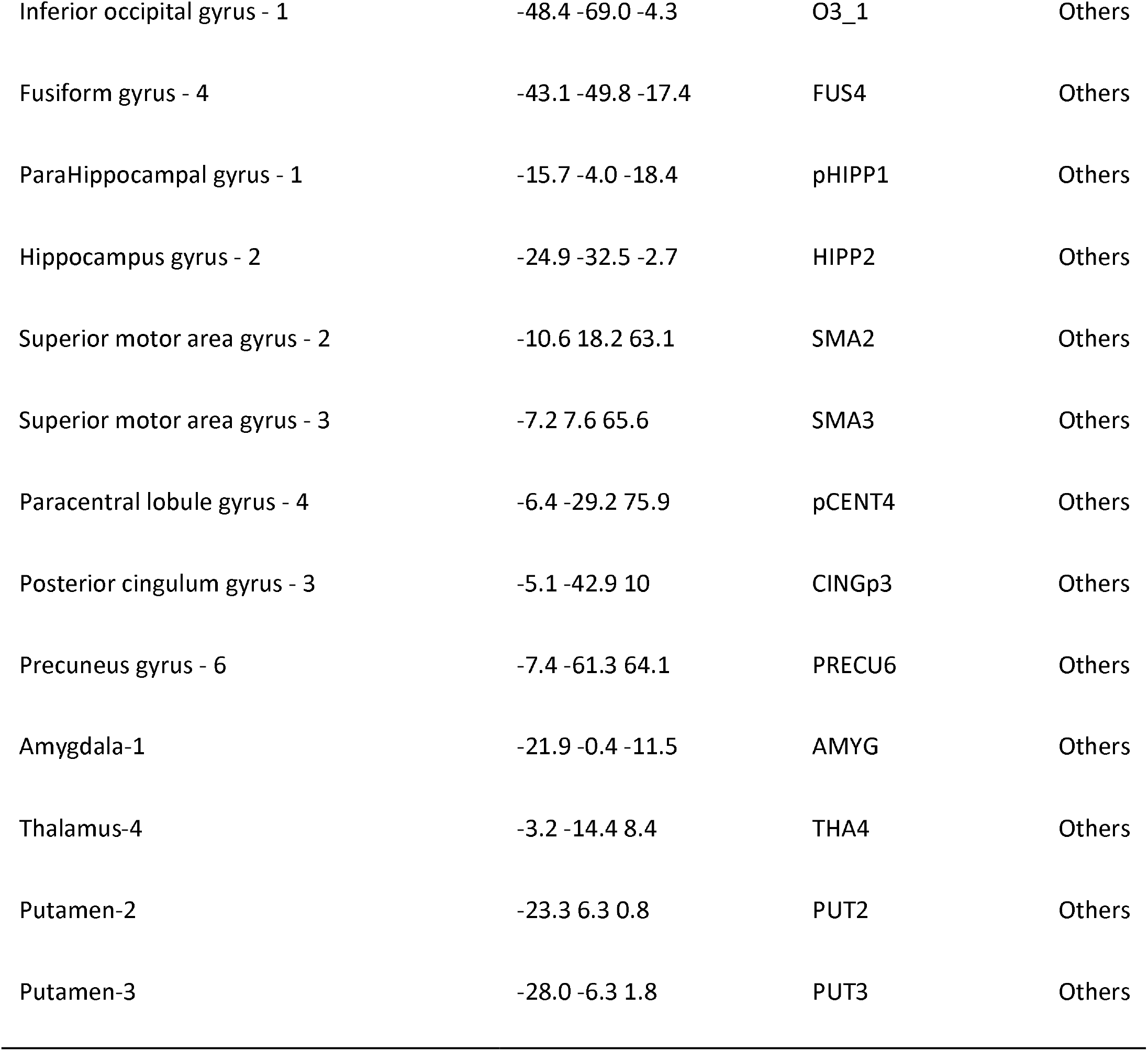
Left hemisphere nodes.

#### 2.4.2 Functional Connectivity

Functional connectivity was computed between the different regions of interest (ROI) using the nodes from previous task-based fMRI work by Labache et al. (Labache et al., 2019) to identify nodes in the dorsal and ventral streams with the peak coordinates from the two tasks used by Labache et al. The nodes were defined as 6mm radius spheres around the peak coordinates. To better fit the dual-stream model’s bilateral ventral streams, we also included the homologue of temporal and parietal nodes in the right hemisphere to study the activity in the right ventral stream. We then calculated the Pearson correlation coefficients between each pair of nodes; correlation coefficients were then Fisher transformed. To explore possible functional reorganization in the right hemisphere in response to left dorsal stream damage, we also included potential right dorsal stream nodes using the homologue coordinates of the left hemisphere’s dorsal stream’s nodes. All the functional connectivity analyses were performed using in-house scripts executed in MATLAB.

##### 2.4.2.1 Defining the Dual-Stream Speech Processing Network in Control Subjects

To confirm that the neural correlates of the dual-stream model do in fact form a significant network in our neurotypical control participants, we computed the following paired-samples t-tests within the control group : 1) Comparison of the average functional connectivity within the entire, bilateral dual-stream network, as well as the average connectivity in just the left hemisphere nodes of the dual-stream network (i.e., between and within the left dorsal and left ventral streams), with that of within the well-defined default mode and visual networks, and 2) Comparison of the average functional connectivity between each stream of the dual-stream network (i.e. bilateral ventral, and left dorsal) to the average functional connectivity between the dual-stream nodes and those of the default mode network and visual network. The coordinates of the default mode network nodes (Table 4) used have been well-defined by numerous previous resting-state and task-related fMRI studies (Gao & Lin, 2012; Vincent, Kahn, Snyder, Raichle, & Buckner, 2008; H. Zhu et al., 2016). The visual network coordinates (Table 4) were taken from a previous task-based fMRI study (Gao & Lin, 2012), which aligns well with numerous other works identifying this network (Lee, Smyser, & Shimony, 2013; Schöpf et al., 2010; Tedeschi et al., 2016; Van Den Heuvel & Pol, 2010). A false discovery rate (FDR) correction was used to control the false positives at *p*<0.05. All the statistical analyses including paired t-tests, two-sample t-tests, regression models and multiple comparison corrections were coded in R.

**Table 4.** Default mode network and visual network nodes examined for comparison purposes.

| Region | MNI Coordinates | Category |
| --- | --- | --- |
| Left hippocampus formation | -21, -15, -14 | Default mode |
| Right hippocampus formation | 24, -19, -21 | Default mode |
| Medial prefrontal cortex | 0, 51, -7 | Default mode |
| Posterior cingulate cortex | 1, -55, 17 | Default mode |
| Left posterior inferior parietal lobule | -47, -71, 29 | Default mode |
| Right posterior inferior parietal lobule | 50, -64, 27 | Default mode |
| Left calcarine | -8, -72, 4 | Visual |
| Right calcarine | 16, -67, 5 | Visual |
| Left cuneus | -5, -96, 12 | Visual |
| Right cuneus | 18, -96, 12 | Visual |
| Left lateral occipital | -23, -89, 12 | Visual |
| Right lateral occipital | 37, -85, 13 | Visual |

##### 2.4.2.2 Functional Connectivity of the dual-stream language network in the stroke group

For the stroke group (in the same fashion as for the control group, above), we computed the overall connectivity of the dual stream model network, as well as the left-hemisphere streams only, and each stream separately, and compared them to that of the visual and default mode networks using paired-samples t-tests with false discovery rate correction to control false positives at *p*<.05.

##### 2.4.2.3 Functional Connectivity comparisons between control and stroke groups

Then, to compare the functional connectivity of the dual-stream nodes of the stroke group to those of the control group, we computed independent-samples t-tests of the average functional connectivities between the stroke and control groups for the following nodes: within left ventral, within left dorsal, between left ventral and left dorsal, between right dorsal and ventral, within right dorsal, within right ventral, between left and right ventral, between left and right dorsal, and across left ventral, left dorsal and right ventral. An FDR correction was used to control the false positives at *p*<0.05.

#### 2.4.3 Graph Theory analysis

Using the functional connectivity correlations described above as the edges between the nodes in table 3 and the homologue in the right hemisphere, we used Graph Theory methods to extract the average shortest path lengths for each node in order to identify hub regions. The parameters for the graph construction are concluded in Table 5 (Achard et al., 2006; M. D. Humphries, Gurney, & Prescott, 2006). To obtain the average shortest path length, first a binary undirected graph at varying thresholds of the functional connectivity was constructed, to identify an appropriate threshold for the subsequent analyses using small-world properties. To construct a small-world network, the graph needs to satisfy ratio γ = *C_net_*/*C_ran_* > 1 and λ = *L_net_*/*L_ran_* ∼ 1 (Achard et al., 2006; Montoya & Solé, 2002; Watts & Strogatz, 1998) and “small-worldness” ratio σ > 1 (Achard et al., 2006; M. D. Humphries et al., 2006). To compute *C_ran_* and *L_ran_*, we applied randomized manipulation by a Markov-chain algorithm (Y. Liu et al., 2008; Maslov & Sneppen, 2002) 100 times for each potential threshold and obtained *C_ran_* and *L_ran_* from the mean of 100 random networks. Then, we found an optimized threshold to attain a best balance between γ, λ, and σ (γ > 1, λ ∼ 1 and σ > 1) using a search of the proportion of possible connections that actually exist between all the nodes. The beginning of the search was a proportion transformation from lowest edges constraint of a small-world network, K = log(N). The end of the search is the 20%, which was the maximum threshold to account for the known sparsity of small-world organization (Achard & Bullmore, 2007; Roger et al., 2019).

**Table 5.**
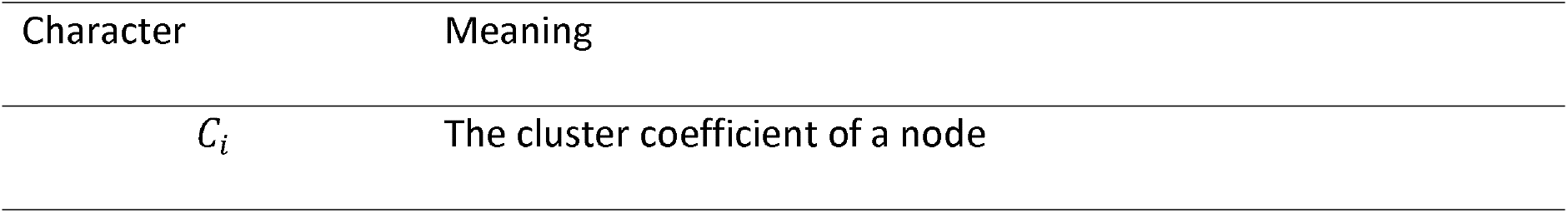
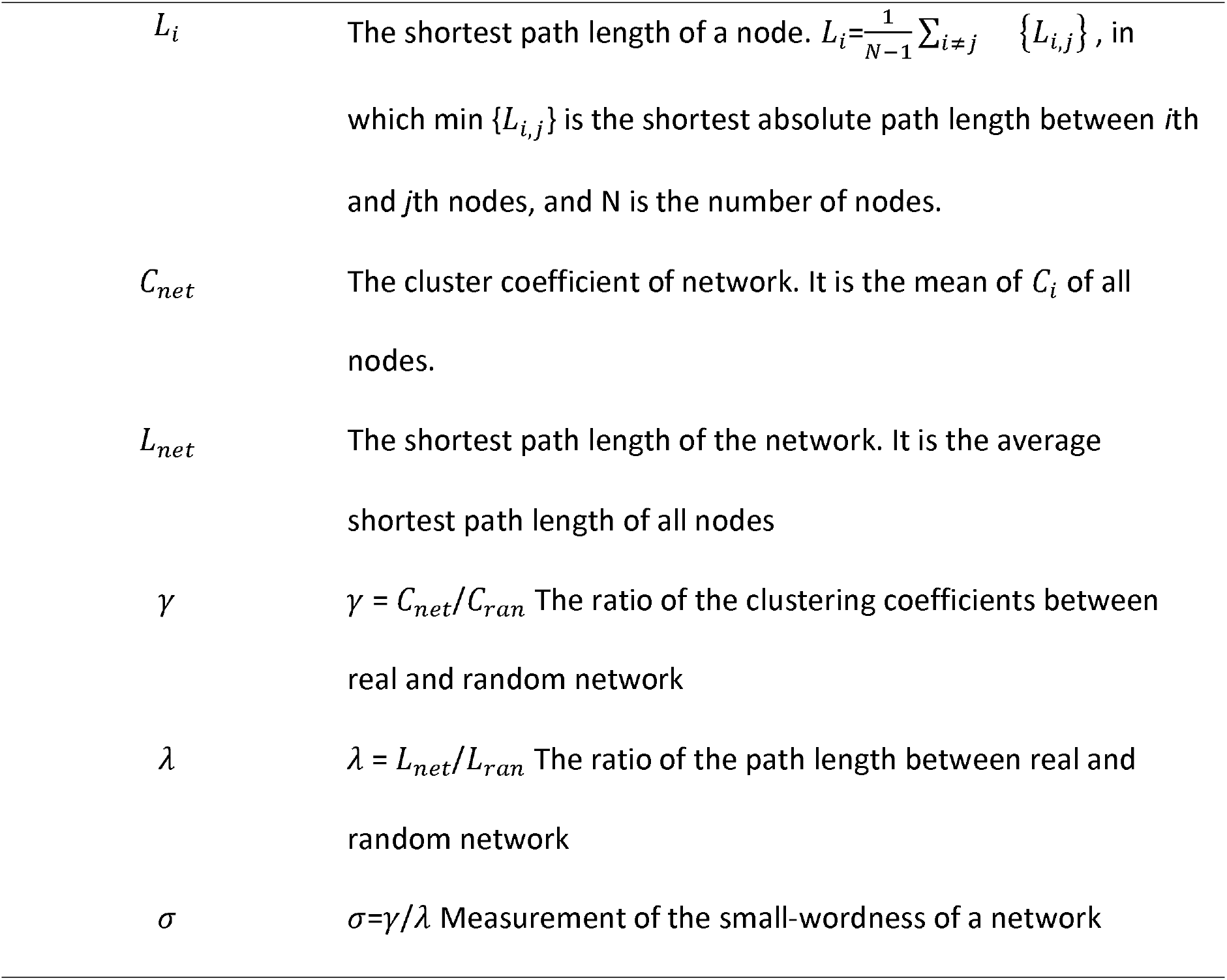
The measurements and their meaning in the brain functional network (Y. Liu et al., 2008)

| Character | Meaning |
| --- | --- |
| $C_i$ | The cluster coefficient of a node |
| $L_i$ | The shortest path length of a node. $L_i = \frac{1}{N-1} \sum_{i \neq j} \{L_{i,j}\}$, in which $\min \{L_{i,j}\}$ is the shortest absolute path length between $i$th and $j$th nodes, and $N$ is the number of nodes. |
| $C_{net}$ | The cluster coefficient of network. It is the mean of $C_i$ of all nodes. |
| $L_{net}$ | The shortest path length of the network. It is the average shortest path length of all nodes |
| $\gamma$ | $\gamma = C_{net}/C_{ran}$ The ratio of the clustering coefficients between real and random network |
| $\lambda$ | $\lambda = L_{net}/L_{ran}$ The ratio of the path length between real and random network |
| $\sigma$ | $\sigma = \gamma/\lambda$ Measurement of the small-worldness of a network |

We then identified “hub” nodes based on the mean shortest path lengths computed. Again, the mean shortest path length formula of a node is: 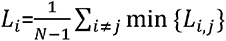. To be identified as a hub of the dual-stream network, a node must satisfy two criteria: 1) be one of the two nodes with smallest *L_i_* values in left dorsal, right dorsal, left ventral, or right ventral streams; 2) be a node with functional connectivity to at least half of the other nodes either directly or mediated by one or more nodes that passes the threshold defined above. Then, to continue our exploration of a possible right dorsal stream involvement in aphasia, we labeled the homologues of the hubs in the left dorsal stream as hubs. Independent-samples t-tests then were used to detect the functional connectivity differences within and between the streams’ hub nodes in and between the control and stroke groups. Once again, the significant results were FDR-corrected to p<0.05.

#### 2.4.4 Regression Analysis of Hub Node Functional Connectivity to Predict Stroke Group Performance

The hub nodes defined above in the control group were used to calculate the functional connectivity between hubs nodes in the stroke group. Then, the functional connectivities of the hub nodes were compared between groups using an independent-samples t-test. The functional connectivities within each stream also were used as predictors in multiple regression models (*lm* command in R) that were applied to predict each stroke survivor’s performance on each WAB behavioral measure. The selection of the predictors included to predict each measure from the WAB, were based on previous lesion-symptom mapping and task-based fMRI work (Baldo, Arévalo, Patterson, & Dronkers, 2013; Fridriksson et al., 2018; Kertesz, 2022; Kümmerer et al., 2013; Shulman et al., 1997; Thye & Mirman, 2018), and are summarized in Table 6. Lesion size, age, gender and education years were also included in each model as covariates. We also applied a reduction of predictor variables using Akaike Information Criterion based elimination in R (step order) to remove the predictors or covariates that contribute little to each model – a standard procedure to remove predictors that are not related to the response variable that could lead to an inflated prediction error (Chambers & Hastie, 1992).

**Table 6.** The functional connectivity predictors for WAB-R.

|  | L Dorsal | R Dorsal | L Ventral | R Ventral | L<br>Dorsal-Ventral | R<br>Dorsal-Ventral | L & R<br>Dorsal | L & R<br>Ventral |
| --- | --- | --- | --- | --- | --- | --- | --- | --- |
| Spont. Speech | ? | ? |  |  | ? | ? | ? |  |
| Auditory Verbal<br>Comprehension |  |  | ? | ? | ? | ? |  | ? |
| Repetition | ? | ? | ? | ? | ? | ? | ? | ? |
| Naming and<br>word finding | ? | ? | ? | ? | ? | ? | ? | ? |
| Aphasia<br>quotient | ? | ? | ? | ? | ? | ? | ? | ? |

## Results

### 3.1 Functional connectivity of the dual stream model

*Control Group.* The paired-samples *t*-tests computed within the control group to compare the mean functional connectivity within the dual stream network with that of the other reference networks yielded the following results: the functional connectivity of the visual network was significantly higher than that of the dual-stream network (t(27)=7.97, FDR p<0.05), the default mode network (t(27)= 5.97, FDR p<0.05), and the left hemisphere dual-stream network (t(27)=7.65, FDR p<0.05; Figure 2). There was no significant difference between the functional connectivity of the dual-stream and default mode networks and the left hemisphere dual-stream.

**Figure 1.**
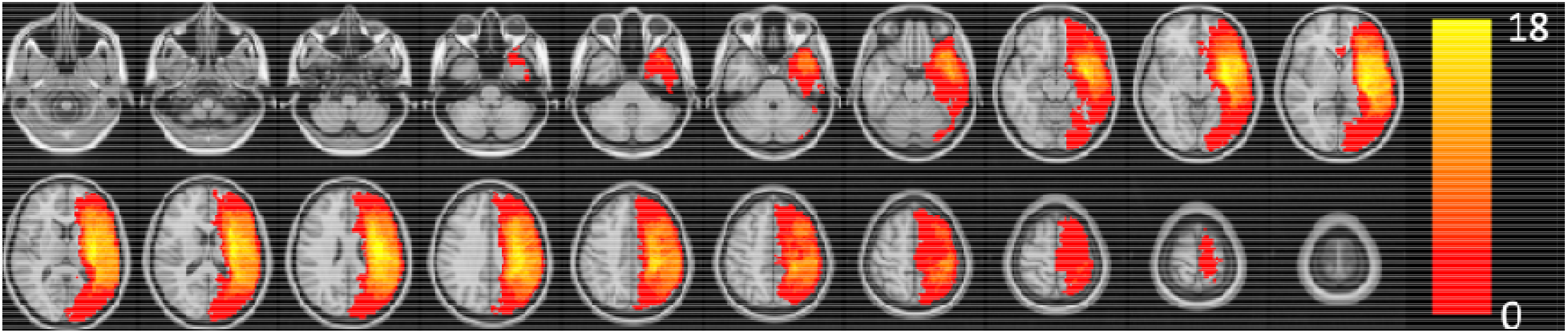
Lesion overlap map of stroke group.

**Figure 2.**
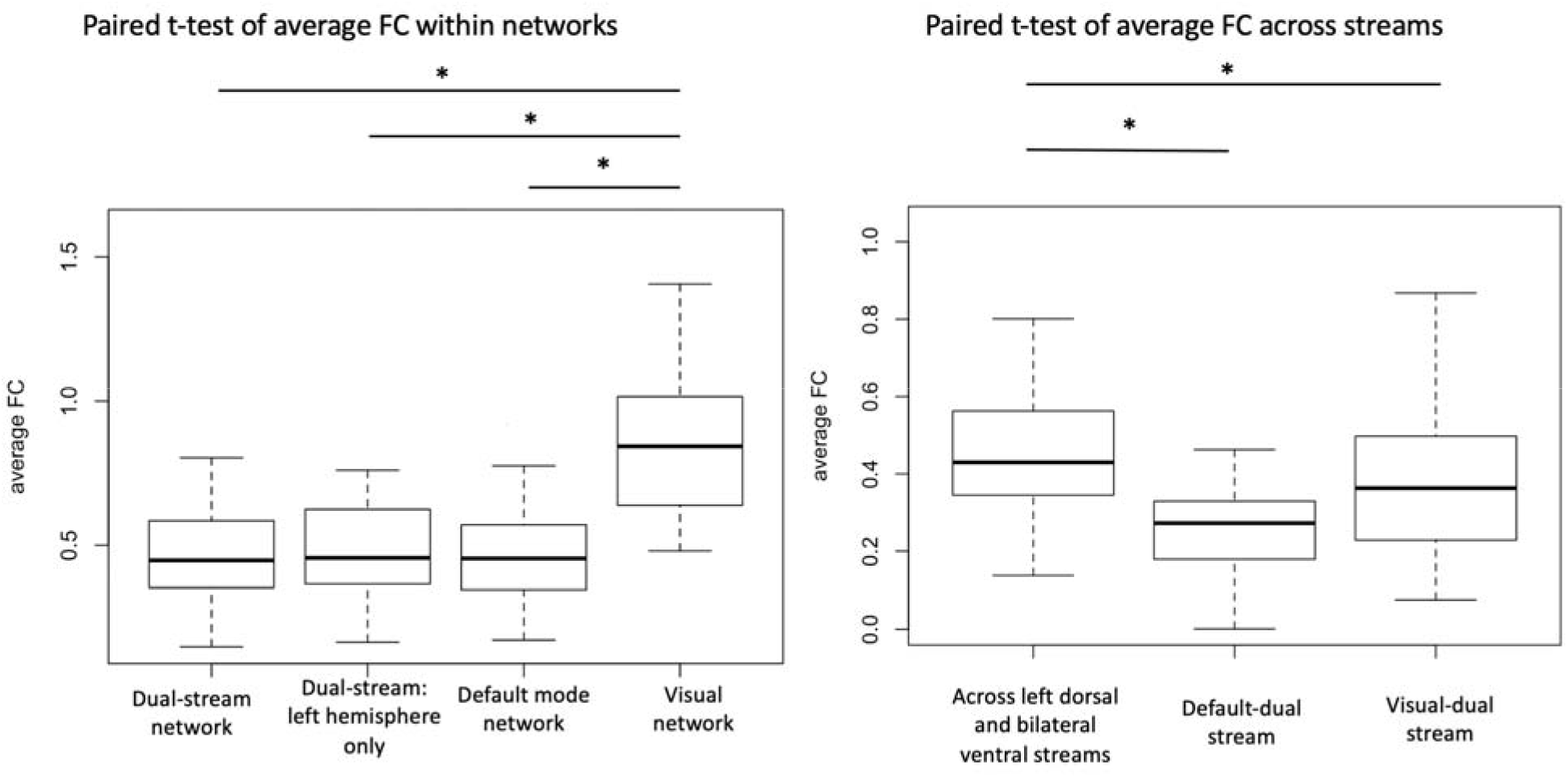
Control group network comparisons. Left: comparison of the functional connectivity within dual stream network, within left hemisphere (between and within left dorsal and ventral stream), default mode network, and visual network. Right: comparison of the functional connectivity within right dorsal and bilateral ventral streams. Asterisk indicates statistical significance at *p*<0.05, FDR corrected.

*Stroke Group.* Similar to the control group, the stroke group exhibited functional connectivity within the visual network that was higher than that of the dual-stream network (t(27)=6.00, FDR p<0.05), the default mode network (t(27)=4.61, FDR p<0.05 and the left part of dual-stream network (t(27)=6.16, FDR p<0.05); Fig. 3). No significant result was found between the dual-stream network, default mode network, and left hemisphere dual-stream.

**Figure 3.**
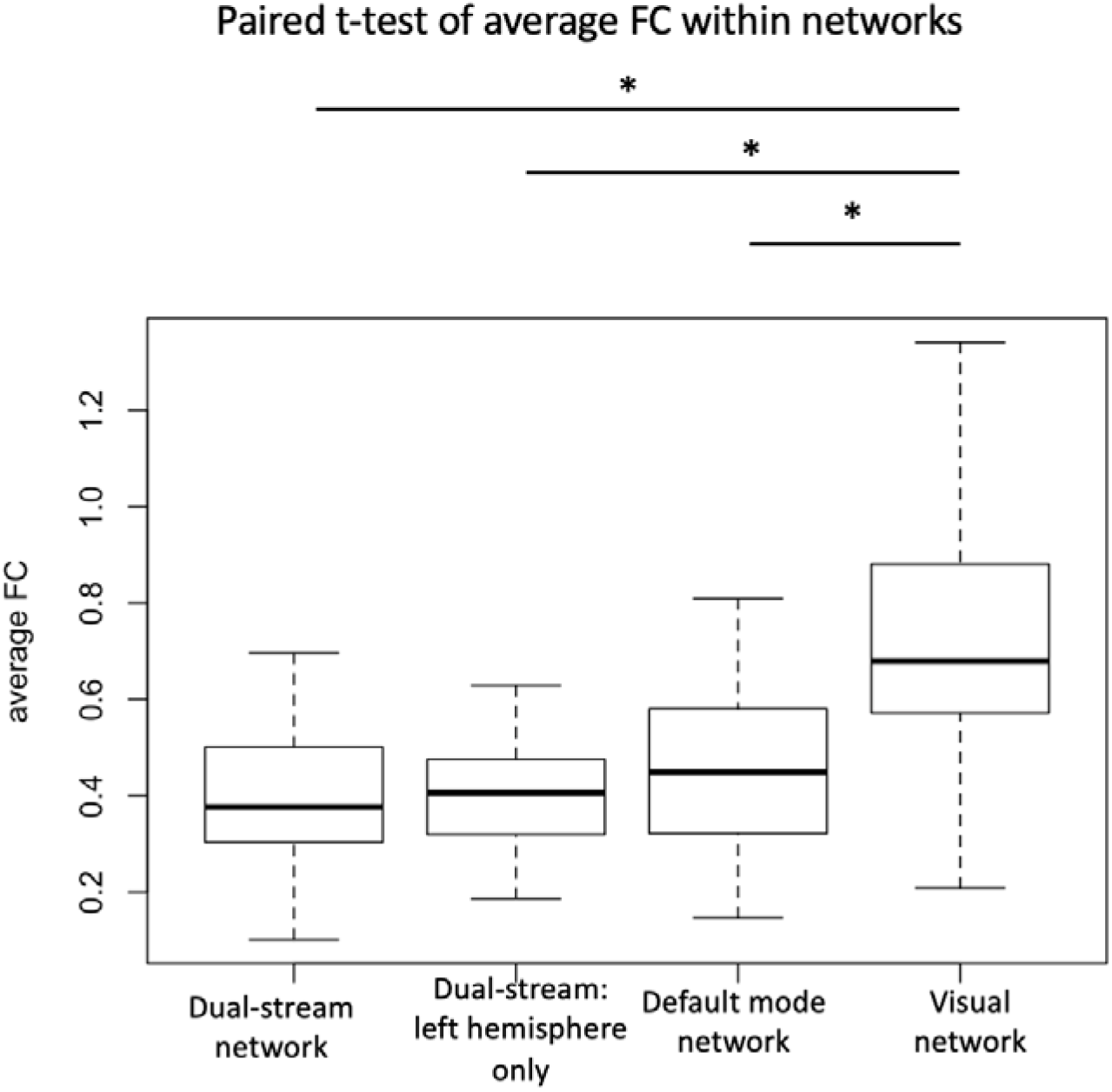
Stroke group network comparisons. Comparison of the functional connectivity within dual stream network, within only the left hemisphere of the dual-stream network (i.e. between and within left dorsal and ventral stream), default mode network, and visual network. An asterisk indicates statistical significance at *p*<0.05, FDR corrected.

**Figure 4.**
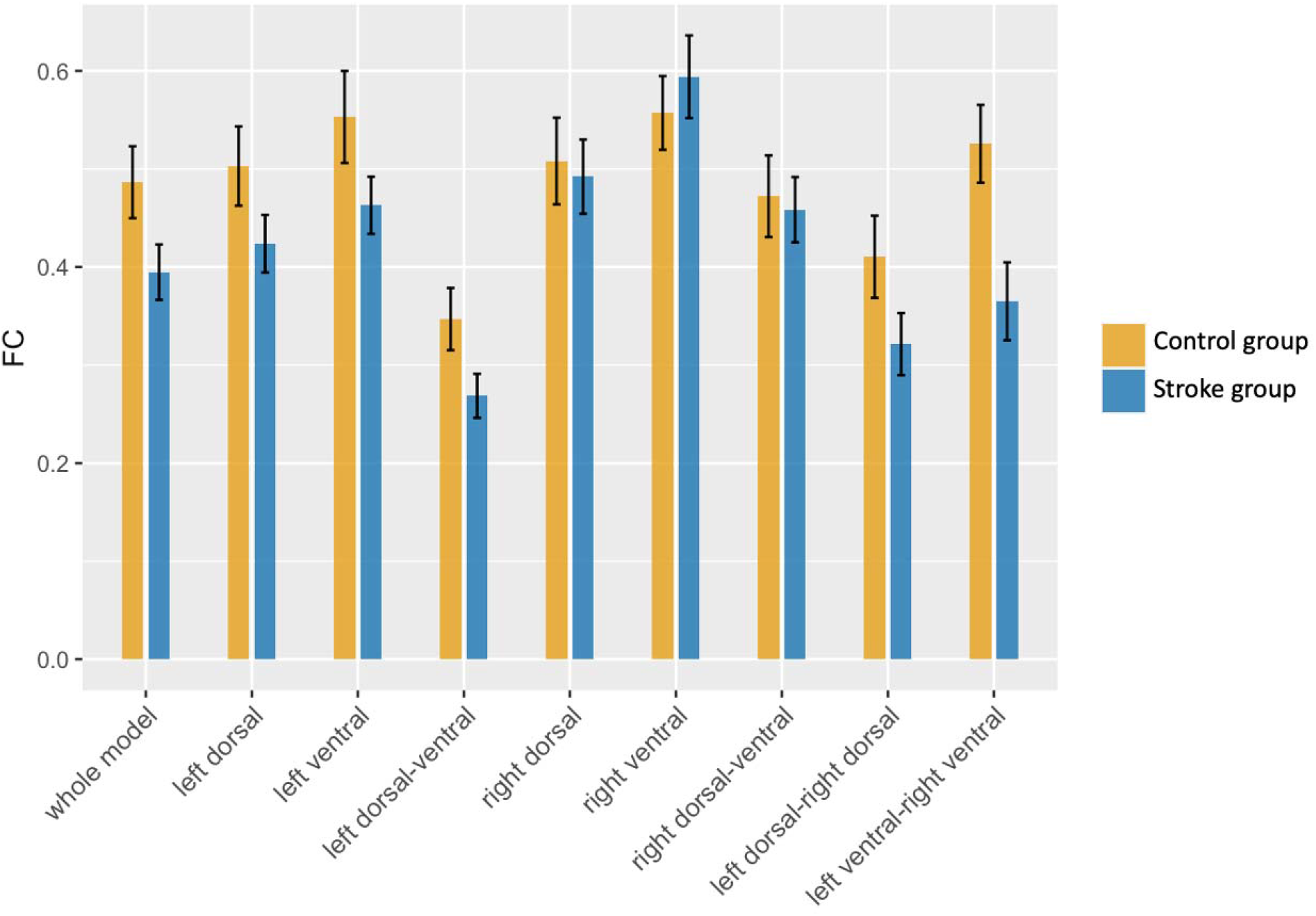
Mean functional connectivity of all nodes within different portions of the dual-stream network for the control and stroke groups. There is no significant result after FDR correction at p<0.05. The error bar indicates one standard deviation.

### 3.2 Functional connectivity across groups in typical dual-stream regions

There was no significant difference of the average functional connectivity in typical dual-stream regions across groups.

### 3.3 Construction of graph with small world network assumption

To construct a graph that satisfies the small world network assumption, we need γ > 1, λ ∼ 1, and σ > 1 (Achard et al., 2006). Fig. 5 indicates the trend of λ, γ, and σ as the proportion of connections is increased. The λ is closest to 1 at proportion 0.132, and the highest value of γ and σ also occur at threshold 0.132. Therefore, the best proportion of connections in our data is 0.132. The small world parameters of all the nodes are shown in Table 7.

**Figure 5.**
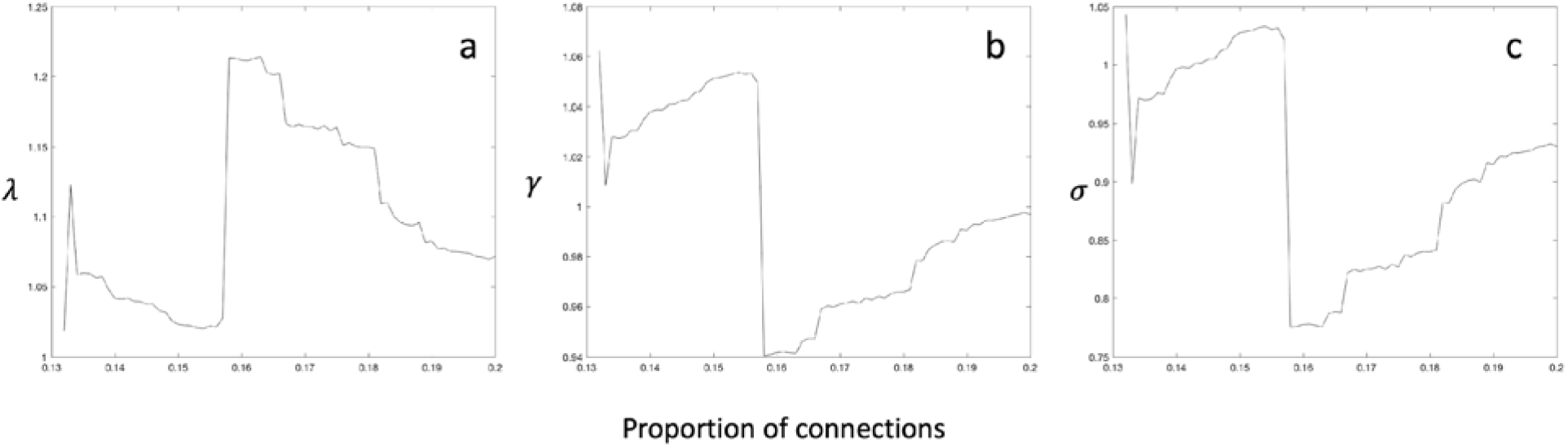
Small-world properties of brain network as a function of proportion of connections. a. λ approaches 1 most at threshold 0.132. b. γ has the highest value at threshold 0.132; c. σ has the highest value at threshold 0.132.

**Table 7.** The small-world network parameters

| Proportion threshold | $L_{net}$ | $C_{net}$ | $\lambda$ | $\gamma$ | $\sigma$ |
| --- | --- | --- | --- | --- | --- |
| 0.132 | 2.26 | 0.54 | 1.02 | 1.06 | 1.04 |

### 3.4 Hubs of the dual-stream network

Eight nodes of the dual-stream network were identified as hubs (i.e., defined by the top two ranked shortest path length nodes in each stream in left dorsal and bilateral ventral, right ventral hubs are the homologue of the left side). The hubs identified were the following nodes: left superior temporal sulcus and left pars triangularis in the left dorsal stream, the homologues in the right dorsal, middle portion of the left superior temporal gyrus and posterior portion of the left middle temporal gyrus for the left ventral stream, and the middle portion of the right middle temporal gyrus and the middle portion of the right superior temporal sulcus of the right ventral stream (Table 8 and Fig 6).

**Figure 6.**
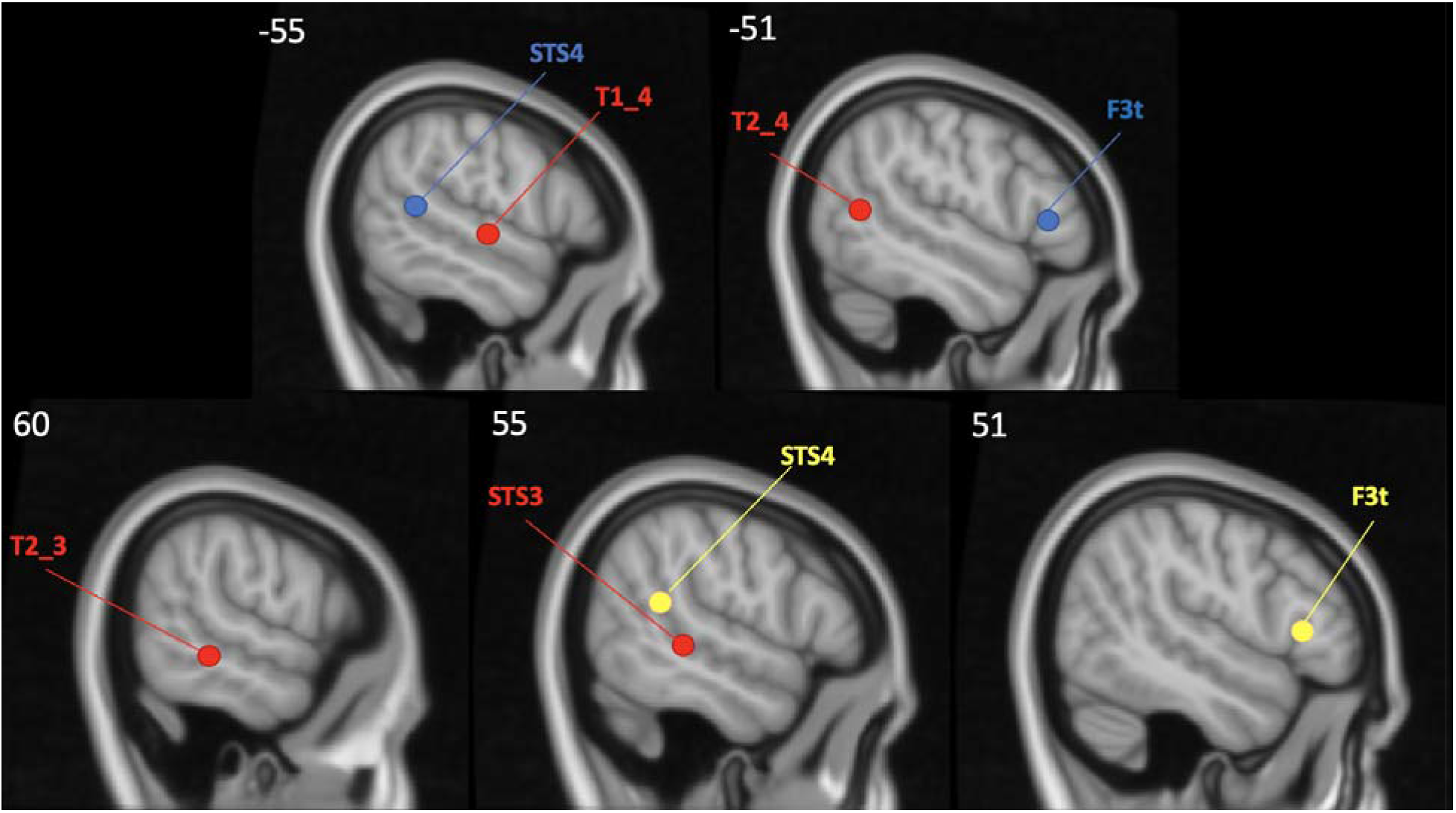
The hub nodes of the dual-stream language network identified in the control group. Red indicates the ventral stream, blue indicates the left dorsal stream, yellow indicates the right dorsal stream which is the homologue of the left dorsal stream. The top row indicates the hub nodes identified in the left hemisphere, and the bottom row indicates nodes identified in the right hemisphere. The caption of the nodes can be found in Table 3.

**Table 8.**
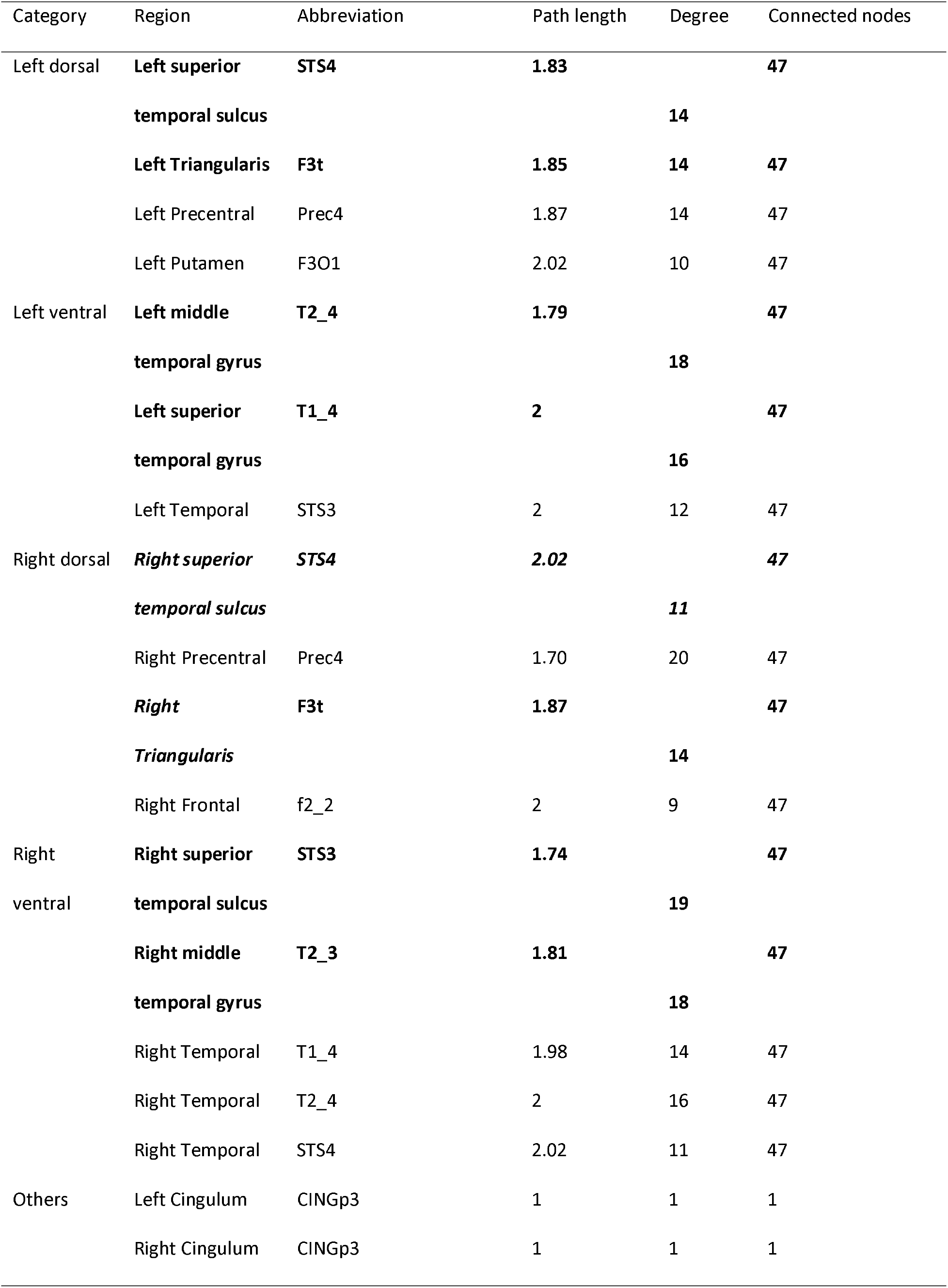
Properties of the top ranked lower average path length nodes in bilateral dorsal and ventral stream. Bold indicates the hub nodes, and the italics indicate the right hemisphere homologues of the left dorsal hubs. Note that if we were to find the lowest average path length nodes as the hub in the right dorsal stream directly, the nodes identified remain the same.

### 3.5 Functional connectivity between hubs in the control and stroke groups

Independent t-tests examining group functional connectivity differences of hub regions, within or between dorsal and ventral streams, resulted in the following significant results (Fig 6): the functional connectivity within the left dorsal stream of the control group was significantly higher than that of the stroke group (t(54)=2.84, FDR p<0.05) and the functional connectivity between the left ventral and right ventral streams of the control group was significantly higher than that of the stroke group(t(54)=2.30, FDR p<0.05).

### 3.6 Regression model to predict WAB performance with hub functional connectivity

Table 9 lists all of the results from the regression models predicting each WAB measure from the functional connectivities of the dual-stream network’s hubs in the stroke group. Significant results were as follows: Spontaneous Speech’s significant predictors were left dorsal (*β*=-15.44, t=-3.51, p<0.01) and right dorsal-ventral (*β*=-7.21, t=-2.21, p<0.05) functional connectivity, as well as age (*β*=-0.19, t=-2.64, p<0.05), education (*β*=0.78, t=2.36, p<0.05) and lesion size (*β*=-1.0e-04, t=-2.28, p<0.05). Right dorsal-ventral functional connectivity (*β*=-3.17, t=-2.13, p<0.05) and age (*β*=-4.0e-05, t=-2.54, p<0.05) were the only significant predictors of the Auditory Verbal Comprehension score. Repetition scores were significantly predicted by left dorsal (*β*=-4.92, t=-2.92, p<0.01), bilateral dorsal (*β*=6.55, t=2.58, p<0.05), right ventral (*β*=2.11, t=2.44, p<0.05), and right dorsal-ventral (*β*=-5.37, t=-2.84, p<0.05) functional connectivity, as well as education (*β*=0.34, t=2.20, p<0.05). Naming and Word Finding scores were significantly predicted by left dorsal (*β*=-5.38, t=-3.08, p<0.01), bilateral dorsal (*β*=5.38, t=2.11, p<0.05), right ventral (*β*=2.80, t=3.45, p<0.01), and right dorsal-ventral (*β*=-6.88, t=-3.75, p<0.01) functional connectivity, as well as education (*β*=0.37, t=2.34, p<0.05). Aphasia Quotient yielded the following significant predictors: left dorsal (*β*=-56.83, t=-3.13, p<0.01), bilateral dorsal (*β*=58.76, t=2.93, p<0.01) and right dorsal-ventral (*β*=-41.74, t=-2.67, p<0.05) functional connectivity, as well as education (*β*=3.76, t=2.88, p<0.01).

**Table 9.**
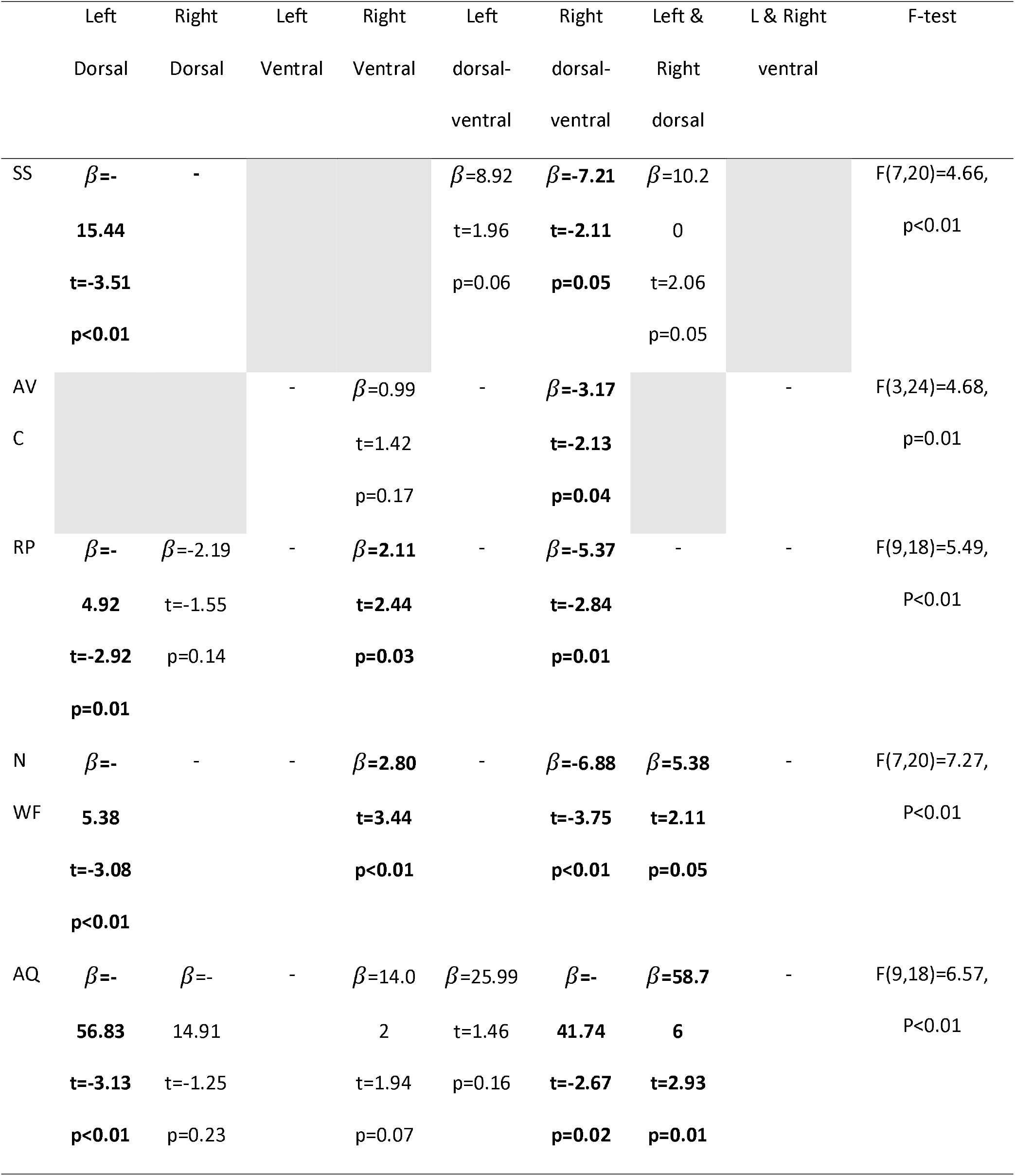
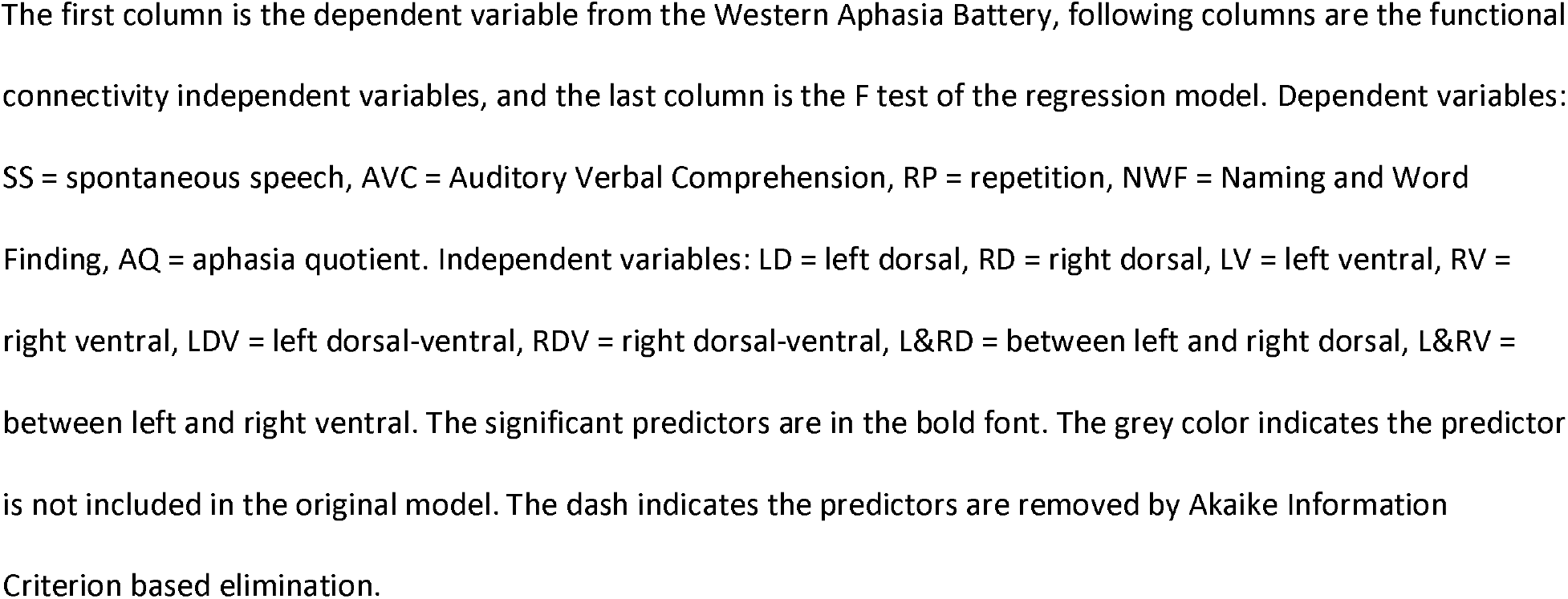
The regression model using the hub functional connectivity predictors

## Discussion

The purpose of this study was two-fold: (1) characterize the dual-stream model’s network characteristics in a neurotypical control group and (2) determine the feasibility of using these dual-stream network characteristics to predict speech and language impairments in stroke survivors. Our findings indicated that: (1) the regions within the dual-stream model of speech processing do perform as an intrinsic network in resting-state fMRI; (2) stroke survivors exhibited significantly less functional connectivity of the hub regions identified by the graph theory measures compared to the neurotypical control group, and (3) functional connectivity of hub regions, but not overall functional connectivity or efficiency measures are significant predictors of post-stroke language performance.

### 4.1 Dual-stream model represents an intrinsic resting-state network

In previous studies, the intrinsic connectivity of a network is defined as ongoing neural and metabolic activity that occurs in the resting basal state (Kilpatrick et al., 2011; Napadow et al., 2010; Raichle, 2015). After the default mode network was discovered in the resting-state (Raichle et al., 2001), many other functional networks were studied with resting-state fMRI, including but not limited to executive, salience, dorsal attention, and visual networks (Napadow et al., 2010; Tessitore et al., 2017; H. Zhu et al., 2016). To test if the regions identified by the dual-stream model of speech processing also can be treated as an intrinsic functional network at rest, we computed two sets of comparisons: 1) the functional connectivity within the dual-stream network compared to functional connectivity within two well-known networks – the default mode network and the visual network, and 2) the functional connectivity between each of the three main components of the dual-stream model (i.e., left dorsal, left ventral, and right ventral) compared to between the dual-stream, default-mode, and visual networks. Our results indicated no significant difference between the strength of the functional connectivity within the default mode and within the dual-stream network, but the visual network exhibited significantly higher functional connectivity than the default mode and dual-stream network (Fig. 2). It is not surprising that the visual network had the strongest functional connectivity, because the locations of the visual nodes are spatially very close to one another compared to the other two networks. While null results should be interpreted cautiously, the non-significant results between the default mode and dual-stream networks indicates that the strength of connections within the dual-stream network are similar to those of the default mode network. This finding suggests the intensity of intrinsic activity within the dual-stream regions is similar to the most well-known resting-state network, which provides robust evidence for the existence of an intrinsic language network as outlined by the dual-stream model of speech processing. In addition, the connections between each of the streams were significantly stronger than the connections between the streams and the visual and default-mode networks. If the dorsal and ventral streams do not belong to the same intrinsic network, these between stream connections would be the same or even weaker than the functional connectivity between the dual-stream regions and regions in these other networks. Based on these findings, we highly recommend that the dual-stream model can be interpreted as an intrinsic network in neurotypical adults, providing further evidence that there is potential to use resting-state functional connectivity measures of the dual-stream network to study the integrity of language circuitry in stroke survivors (Battistella et al., 2020). Our resting-state functional connectivity findings are highly consistent with previous task-based functional MRI studies that have shown that regions in the dual-stream model are co-activated during a variety of language tasks (Hickok & Poeppel, 2004; Saur et al., 2008; Saur et al., 2010).

To further characterize the dual stream network, we identified hub regions of the resting-state dual-stream network using graph theoretical parameters. The hub regions identified in the current study were superior temporal sulcus and triangularis for the dorsal stream, middle temporal gyrus and superior temporal gyrus for left ventral stream, as well as superior temporal gyrus and middle temporal gyrus for right ventral stream. These regions are all implicated in numerous lesion-symptom mapping and neuroimaging studies (Dronkers, 2011; Margulies & Petrides, 2013; Mesulam et al., 2014; Price, 2012; Vigneau et al., 2006). The left pars triangularis, i.e. the anterior portion of Broca’s area, is reliably implicated in speech production, including naming and phonological awareness (Foundas, Eure, Luevano, & Weinberger, 1998; Foundas, Leonard, Gilmore, Fennell, & Heilman, 1996; Kibby, Kroese, Krebbs, Hill, & Hynd, 2009). The other hub node in the dorsal stream is near the superior temporal sulcus, which is also known as the audiovisual integration area (Heim et al., 2008). The ventral stream hubs include bilateral nodes in the mid and posterior middle temporal gyrus, overlapping with bilateral middle temporal regions that are reliably implicated in speech comprehension, particularly lexical-semantic processing (Acheson & Hagoort, 2013; Wallentin et al., 2011). Lesion-symptom mapping studies of stroke consistently identify damage in our left hemisphere middle temporal hubs as associated with auditory comprehension deficits (Bates et al., 2003; Dronkers, Wilkins, Van Valin Jr, Redfern, & Jaeger, 2004; Hickok & Poeppel, 2007; Pillay, Binder, Humphries, Gross, & Book, 2017, Rogalsky et al., 2022;). Thus, the hub nodes identified by our graph theory approach are not surprising, but it is noteworthy that the right ventral hubs are in the middle temporal gyrus and not the superior temporal gyrus. Most task-based neuroimaging and neurostimulation studies of speech identify more superior temporal than middle temporal regions in the right hemisphere, which is often attributed to spectrotemporal acoustic processing that is not specific to speech (Hickok & Poeppel, 2007; Wolmetz, Poeppel, & Rapp, 2011). Our findings suggest that multiple regions of the right middle temporal gyrus are highly connected with other nodes of the dual-stream network, and require further study as to their role in speech processing and aphasia recovery.

### 4.2 Functional connectivity patterns in stroke survivors

Numerous previous studies that have employed lesion-symptom mapping indicate that structural damage to the dual-stream regions is related to various speech and language impairments in post-stroke aphasia (Agosta et al., 2013; Kümmerer et al., 2013; Yang et al., 2017). However, studies of functional differences within and across the dual streams, beyond the area of infarct, are a promising direction but relatively rare (Battistella et al., 2020). Connectome-based lesion-symptom mapping is one way to characterize disruptions in the structural integrity of white matter tracts connected to, but beyond the area of clear, structural infarct (Baboyan et al., 2021; Matchin et al., 2022). This technique entails demarcating the area of lesion and using that area as a seed region to identify, in control subjects, white matter tracts and associated networks that are impacted by the focal area of lesion. We see our study using functional connectivity as complementary to the structural network findings from connectome-based lesion-symptom mapping. Our between-group comparisons provide a glance at how intrinsic functional connections may differ, on average, between control participants and individuals post-left hemisphere stroke. We calculated two sets of comparisons between the control and stroke survivor groups: 1) the average functional connectivity across all nodes in each stream of the dual-stream network (Fig. 4), and 2) the average stream functional connectivity across just the hubs nodes of each stream (Fig. 7), identified by the graph theory procedures. No significant differences were observed between the groups when examining the functional connectivity of all nodes in each stream. However, the hub nodes’ functional connectivity comparisons did find significantly weaker connectivity amongst the hub regions of the left dorsal stream, as well as amongst the hub regions of the bilateral ventral stream, in the stroke group compared to the control group. Based on these group-level comparisons, we conclude that graph theory approaches provide meaningful information regarding network-level dysfunction post-stroke, and that these dual-stream hubs defined by graph theory should be investigated further for possible sites of neurostimulation to promote language rehabilitation in stroke survivors. Given the fact that the majority of our stroke group has a diagnosis of aphasia (n = 25 of 28), it is likely that weakened hub node functional connectivity, versus overall node connectivity in the dual-stream network, is strongly related to an aphasia diagnosis post left hemisphere stroke, although future studies are needed to explore this further.

**Figure 7.**
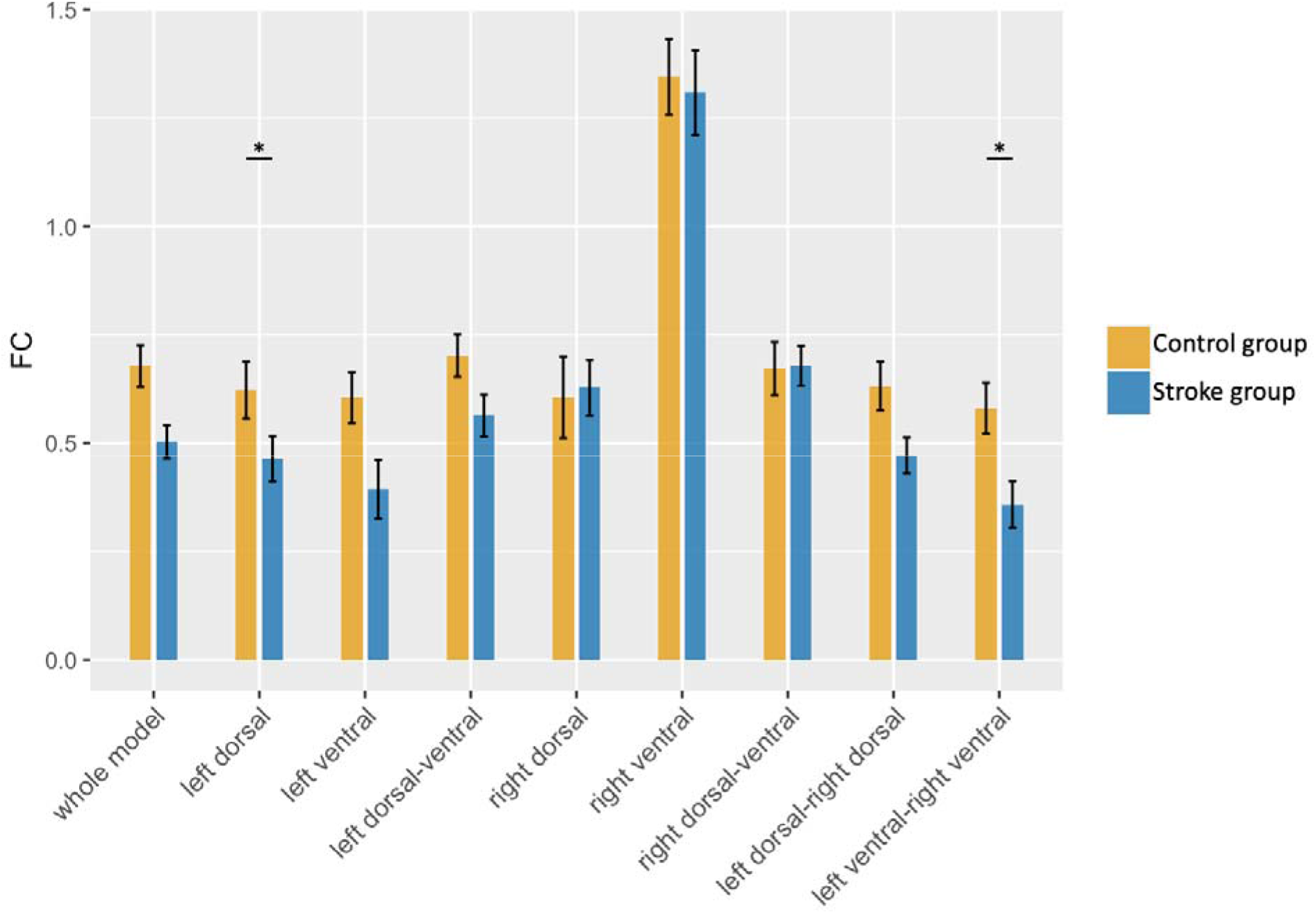
The between-group comparisons of functional connectivity of hubs across the control and stroke groups. Asterisk indicates a significant difference, p<0.05, after FDR correction. The error bar indicates one standard deviation.

### 4.3 Dual-stream functional connectivity as a predictor of language abilities

Due to the significantly lower average functional connectivity between the hub nodes in the stroke group compared to the control group, we tested the ability of these hub functional connectivities to predict performance on subtests of the Western Aphasia Battery - Revised, after accounting for variance due to lesion size, age, gender, and education years (Table 8).

In the ventral stream, the hubs were identified to be bilaterally in the posterior middle temporal gyrus and superior temporal sulcus. Notably, neither the functional connectivity of the hub nodes within the left ventral stream nor the functional connectivity of the left ventral hub nodes with other streams’ hubs were significant predictor of performance on any WAB subtest, perhaps providing additional evidence of the bilateral nature of the ventral stream; perhaps partial damage to the left ventral stream does not yield a deficit on clinical measures due to the right ventral streams’ contributions (we did expect that major damage to the left ventral nodes would yield comprehension impairments (Bonilha et al., 2017; Robson et al., 2014), but our stroke group contained only 3.5% of individuals with Wernicke’s aphasia, which is often associated with large left temporal lesions; thus the effect did not reach significance at our group level).

It also is noteworthy that our regression models did not find functional connectivity between the left dorsal and ventral streams to be a significant predictor of performance on any WAB subtest, including repetition and naming which both have been previously shown to reliably engage both left frontal and left posterior temporal cortices (Cappa, Sandrini, Rossini, Sosta, & Miniussi, 2002; Trimmel et al., 2018). However, it should be noted that for Spontaneous Speech scores, left dorsal-ventral connectivity approaches significance (p = .052). To our knowledge there is no clear theoretical basis or finding in previous work that could explain these null results, but we will speculate here, in hopes that it may be potentially helpful to direct future work. According to the dual-stream model, the dorsal stream and ventral stream communicate mainly via the auditory-motor interface in the posterior superior temporal gyrus (i.e., area Spt) and the arcuate fasciculus, and also via white matter connections between the anterior temporal lobe and inferior frontal gyrus. These connections in the dual-stream model are supported by numerous structural connectivity studies (Glasser & Rilling, 2008; Rilling et al., 2008; Saur et al., 2008). But notably, our hub nodes identified by our graph theory approach did not include any regions in the anterior temporal lobes. Thus, the typical communications between the two streams may not be reflected in our functional connectivity measures between the dorsal and ventral hubs. Another possibility is that the functional connectivity between the left dorsal and ventral hub nodes may not be necessary for successful language abilities, and perhaps even the synchronization of these hub nodes may suggest network dysfunction that leads to lower performance on some language measures.

Our findings related to the right hemisphere homologue of the left dorsal stream (i.e. “right dorsal”) may provide some clarity as to why previous work investigating potential benefits of the right hemisphere in post-stroke recovery have been equivocal (François et al., 2019; Xing et al., 2016). Notably, across all WAB subtests, greater right dorsal-ventral functional connectivity predicted poorer performance. This finding coincides with previous evidence implying that activity in the right hemisphere following left hemisphere stroke is not helpful to language rehabilitation (Karbe et al., 1998; Richter, Miltner, & Straube, 2008), i.e. seemingly supporting a negative view of the compensation activity in the right hemisphere. However, our results provide a more nuanced view of the right hemisphere’s contributions to post-stroke speech and language abilities: while greater right dorsal-ventral functional connectivity was related to poorer performance and more severe aphasia, greater functional connectivity between the left and right dorsal streams was a significant predictor of more mild aphasia symptoms overall (i.e. a higher WAB aphasia quotient). These findings suggest that right dorsal stream contributions to post-stroke speech could possibly be beneficial, but not by replacing the functionality of the left dorsal stream, but rather supporting the remaining left dorsal stream via more domain-general cognitive resources that have been previously located to right lateral frontal cortex (Reuter et al., 2011; Huang et al., 2022), and have been shown to be activated by a variety of speech and language tasks, particularly as task demands increase (Van Ettinger-Veenstra et al., 2010; Just et al., 1996).

One possible explanation to consider for these right dorsal findings is lesion size – perhaps larger left hemisphere lesions lead to both greater speech impairments and also greater right hemisphere involvement in speech tasks, reflecting in greater right dorsal-ventral connectivity. However, this is not the case in our study: we found lesion size to be a significant predictor of performance on only one WAB subtest, Spontaneous Speech. Thus, future studies are needed to investigate the possibility of how to promote functional connectivity between left and right dorsal hubs, but not between right dorsal and ventral hubs, in individuals with aphasia to maximize this potential compensatory strategy in aphasia.

Previous task-based fMRI studies have found that greater right hemisphere activations predict greater aphasia severity (LaCroix, James, & Rogalsky, 2021; Postman-Caucheteux et al., 2010; Richter, Miltner, & Straube, 2008). Conversely, in the present study, we found that stronger right ventral functional connectivity was a significant predictor of better naming and repetition performance - two functions known to engage the ventral stream in neurotypical control participants (Hickok & Poeppel, 2004; Ross et al., 2010; Wise et al., 2001; Saur et al., 2008). These findings provide further evidence for the bilaterality of the ventral stream of the language network. However, it is also notable that performance on the Auditory Verbal Comprehension subtest and overall aphasia severity (as measured by WAB-R Aphasia Quotient) were not predicted by right ventral connectivity. These differences in the predictive ability of right ventral connectivity across the WAB subtests may be due to the nature of the WAB’s comprehension subtest which includes many stimuli of increasing complexity at the phrasal and sentence-level. The right ventral stream has been found to support single word comprehension and phonological encoding (Gorno-Tempini et al., 2004; Hickok & Poeppel, 2007), and not modulated by sentence structure or morphosyntactic manipulations (Humphries et al., 2006; Noven et al., 2021). While the right ventral connectivity did not predict overall aphasia severity, our findings related to naming and repetition suggest that right ventral connectivity may help support some lexical-semantic and phonological functions in post-stroke aphasia. Future studies of the right ventral stream’s connectivity in relation to other more specific receptive speech measures are needed to characterize how the right ventral stream is contributing to receptive and productive speech abilities in individuals with aphasia.

Lastly, we would like to discuss our consistent finding of functional connectivity within the left dorsal stream being a significant negative predictor of WAB performance. On the surface, this finding was surprising: how could less coherence within the left dorsal stream predict better performance on speech tasks known to be supported by these left dorsal regions (e.g. spontaneous speech and repetition)? Upon further inspection of the data, it is apparent that individuals with aphasia with large left frontal lesions encompassing the left dorsal stream hub nodes are driving this negative correlation between left dorsal hub connectivity and WAB performance. The functional connectivity of these individuals’ injured left dorsal nodes (i.e., nodes overlaps the lesion area) is very high compared to functional connectivity from the intact nodes, due to the lack of fluctuation in the BOLD response of their nodes. In other words, the functional connectivity of their left dorsal nodes is very strongly correlated because all of these nodes exhibit very little to no BOLD response. To our knowledge this paradox has not been addressed in the previous literature of functional connectivity after a stroke, and thus previous characterizations of functional connectivity changes may need to be interpreted with this issue in mind (Liu et al., 2017; Zhu et al., 2014). Perhaps novel approaches need to be developed to determine how to best consider both correlation and amplitude of BOLD fluctuations to maximize the utility of using functional connectivity predictors to better understand the neurobiology of post-stroke speech and language impairments.

### 4.4 Conclusions

In conclusion, our study has five main findings: 1) The dual-stream model is representative of an intrinsic network as measured by resting-state fMRI; 2) The functional connectivity of the hub nodes of the dual-stream network, but not overall network connectivity, is significantly lower in our stroke group than in the control group; 3) The intrinsic connectivity of dorsal and ventral stream hubs identified in neurotypical control participants significantly predict performance on clinical aphasia measures (e.g. WAB subtests) in individuals with aphasia; 4) greater right dorsal connectivity with the left dorsal stream, and less connectivity with the right ventral stream, significantly predicts better performance in expressive and receptive speech tasks, and lower overall aphasia severity; (5) greater functional connectivity within the right ventral stream significantly predicts better naming and repetition performance in individuals with aphasia.

## Acknowledgement

This work is supported by NIH NIDCD: P50 DC014664 (PI: J. Fridriksson) and R01 DC009659 (PI: G. Hickok).

## References

1. Achard, S., & Bullmore, E. (2007). Efficiency and cost of economical brain functional networks. PLoS Comput Biol, 3(2), e17.

2. Achard, S., Salvador, R., Whitcher, B., Suckling, J., & Bullmore, E. (2006). A resilient, low-frequency, small-world human brain functional network with highly connected association cortical hubs. Journal of Neuroscience, 26(1), 63–72.

3. Acheson, D. J., & Hagoort, P. (2013). Stimulating the brain’s language network: syntactic ambiguity resolution after TMS to the inferior frontal gyrus and middle temporal gyrus. Journal of cognitive neuroscience, 25(10), 1664–1677.

4. Agosta, F., Galantucci, S., Canu, E., Cappa, S. F., Magnani, G., Franceschi, M., … Filippi, M. (2013). Disruption of structural connectivity along the dorsal and ventral language pathways in patients with nonfluent and semantic variant primary progressive aphasia: a DT MRI study and a literature review. Brain and language, 127(2), 157–166.

5. Albert, R., Jeong, H., & Barabási, A.-L. (2000). Error and attack tolerance of complex networks. Nature, 406(6794), 378–382.

6. Andersen, S. M., Rapcsak, S. Z., & Beeson, P. M. (2010). Cost function masking during normalization of brains with focal lesions: still a necessity? Neuroimage, 53(1), 78–84.

7. Baboyan, V., Basilakos, A., Yourganov, G., Rorden, C., Bonilha, L., Fridriksson, J., & Hickok, G. (2021). Isolating the white matter circuitry of the dorsal language stream: Connectome-symptom mapping in stroke induced aphasia. Human brain mapping, 42(17), 5689–5702.

8. Baldo, J. V., Arévalo, A., Patterson, J. P., & Dronkers, N. F. (2013). Grey and white matter correlates of picture naming: evidence from a voxel-based lesion analysis of the Boston Naming Test. Cortex, 49(3), 658–667.

9. Barabási, A.-L. (2012). The network takeover. Nature Physics, 8(1), 14–16.

10. Bassett, D. S., & Bullmore, E. (2006). Small-world brain networks. The neuroscientist, 12(6), 512–523.

11. Bates, E., Wilson, S. M., Saygin, A. P., Dick, F., Sereno, M. I., Knight, R. T., & Dronkers, N. F. (2003). Voxel-based lesion–symptom mapping. Nature neuroscience, 6(5), 448–450.

12. Battistella, G., Borghesani, V., Henry, M., Shwe, W., Lauricella, M., Miller, Z., … Brambati, S. M. (2020). Task-free functional language networks: reproducibility and clinical application. Journal of Neuroscience, 40(6), 1311–1320.

13. Belleau, E. L., Ehret, L. E., Hanson, J. L., Brasel, K. J., Larson, C. L., & deRoon-Cassini, T. A. (2020). Amygdala functional connectivity in the acute aftermath of trauma prospectively predicts severity of posttraumatic stress symptoms. Neurobiology of stress, 12, 100217.

14. Binder, J. R., Frost, J. A., Hammeke, T. A., Bellgowan, P., Rao, S. M., & Cox, R. W. (1999). Conceptual processing during the conscious resting state: a functional MRI study. Journal of cognitive neuroscience, 11(1), 80–93.

15. Boccaletti, S., Latora, V., Moreno, Y., Chavez, M., & Hwang, D.-U. (2006). Complex networks: Structure and dynamics. Physics reports, 424(4-5), 175–308.

16. Bohland, J., Kapse, K., & Kiran, S. (2014). Graph analytic characterization of resting state networks in post-stroke Aphasia. In: Academy of Aphasia, Miami, FL.

17. Bonilha, L., Hillis, A. E., Hickok, G., Den Ouden, D. B., Rorden, C., & Fridriksson, J. (2017). Temporal lobe networks supporting the comprehension of spoken words. Brain, 140(9), 2370–2380.

18. Brett, M., Leff, A. P., Rorden, C., & Ashburner, J. (2001). Spatial normalization of brain images with focal lesions using cost function masking. Neuroimage, 14(2), 486–500.

19. Cappa, S. F., Sandrini, M., Rossini, P. M., Sosta, K., & Miniussi, C. (2002). The role of the left frontal lobe in action naming: rTMS evidence. Neurology, 59(5), 720–723.

20. Cordes, D., Haughton, V. M., Arfanakis, K., Wendt, G. J., Turski, P. A., Moritz, C. H., … Meyerand, M. E. (2000). Mapping functionally related regions of brain with functional connectivity MR imaging. American journal of neuroradiology, 21(9), 1636–1644.

21. Crinion, J., & Price, C. J. (2005). Right anterior superior temporal activation predicts auditory sentence comprehension following aphasic stroke. Brain, 128(12), 2858–2871.

22. Dronkers, N. F. (2011). The neural architecture of the language comprehension network: converging evidence from lesion and connectivity analyses. Frontiers in systems neuroscience, 5, 1.

23. Dronkers, N. F., Wilkins, D. P., Van Valin Jr, R. D., Redfern, B. B., & Jaeger, J. J. (2004). Lesion analysis of the brain areas involved in language comprehension. Cognition, 92(1-2), 145–177.

24. Duncan, E. S., & Small, S. L. (2016). Increased modularity of resting state networks supports improved narrative production in aphasia recovery. Brain connectivity, 6(7), 524–529.

25. Ferguson, M. A., Lim, C., Cooke, D., Darby, R. R., Wu, O., Rost, N. S., … Fox, M. D. (2019). A human memory circuit derived from brain lesions causing amnesia. Nature communications, 10(1), 1–9.

26. Fitzhugh, M. C., Hemesath, A., Schaefer, S. Y., Baxter, L. C., & Rogalsky, C. (2019). Functional connectivity of Heschl’s gyrus associated with age-related hearing loss: a resting-state fMRI study. Frontiers in psychology, 10, 2485.

27. Foundas, A. L., Eure, K. F., Luevano, L. F., & Weinberger, D. R. (1998). MRI asymmetries of Broca’s area: the pars triangularis and pars opercularis. Brain and language, 64(3), 282–296.

28. Foundas, A. L., Leonard, C. M., Gilmore, R. L., Fennell, E. B., & Heilman, K. M. (1996). Pars triangularis asymmetry and language dominance. Proceedings of the national academy of Sciences, 93(2), 719–722.

29. François, C., Ripollés, P., Ferreri, L., Muchart, J., Sierpowska, J., Fons, C., … Garcia-Alix, A. (2019). Right structural and functional reorganization in four-year-old children with perinatal arterial ischemic stroke predict language production. Eneuro, 6(4).

30. Fridriksson, J., den Ouden, D.-B., Hillis, A. E., Hickok, G., Rorden, C., Basilakos, A., … Bonilha, L. (2018). Anatomy of aphasia revisited. Brain, 141(3), 848–862.

31. Gao, W., & Lin, W. (2012). Frontal parietal control network regulates the anti-correlated default and dorsal attention networks. Human brain mapping, 33(1), 192–202.

32. Gorno-Tempini, M. L., Dronkers, N. F., Rankin, K. P., Ogar, J. M., Phengrasamy, L., Rosen, H. J., … & Miller, B. L. (2004). Cognition and anatomy in three variants of primary progressive aphasia. Annals of Neurology: Official Journal of the American Neurological Association and the Child Neurology Society, 55(3), 335–346.

33. Glasser, M. F., & Rilling, J. K. (2008). DTI tractography of the human brain’s language pathways. Cerebral Cortex, 18(11), 2471–2482.

34. Hampson, M., Peterson, B. S., Skudlarski, P., Gatenby, J. C., & Gore, J. C. (2002). Detection of functional connectivity using temporal correlations in MR images. Human brain mapping, 15(4), 247–262.

35. He, Y., Chen, Z., & Evans, A. (2008). Structural insights into aberrant topological patterns of large-scale cortical networks in Alzheimer’s disease. Journal of Neuroscience, 28(18), 4756–4766.

36. He, Y., & Evans, A. (2010). Graph theoretical modeling of brain connectivity. Current opinion in neurology, 23(4), 341–350.

37. Heim, S., Eickhoff, S. B., & Amunts, K. (2008). Specialisation in Broca’s region for semantic, phonological, and syntactic fluency? Neuroimage, 40(3), 1362–1368.

38. Hickok, G., & Poeppel, D. (2000). Towards a functional neuroanatomy of speech perception. Trends in cognitive sciences, 4(4), 131–138.

39. Hickok, G., & Poeppel, D. (2004). Dorsal and ventral streams: a framework for understanding aspects of the functional anatomy of language. Cognition, 92(1-2), 67–99.

40. Hickok, G., & Poeppel, D. (2007). The cortical organization of speech processing. Nature reviews neuroscience, 8(5), 393–402.

41. Huang, W., Li, X., Xie, H., Qiao, T., Zheng, Y., Su, L., … & Dou, Z. (2022). Different Cortex Activation and Functional Connectivity in Executive Function Between Young and Elder People During Stroop Test: An fNIRS Study. Frontiers in Aging Neuroscience, 14.

42. Humphries, C., Willard, K., Buchsbaum, B., & Hickok, G. (2001). Role of anterior temporal cortex in auditory sentence comprehension: an fMRI study. Neuroreport, 12(8), 1749–1752.

43. Humphries, C., Binder, J. R., Medler, D. A., & Liebenthal, E. (2006). Syntactic and semantic modulation of neural activity during auditory sentence comprehension. Journal of cognitive neuroscience, 18(4), 665–679.

44. Humphries, M. D., Gurney, K., & Prescott, T. J. (2006). The brainstem reticular formation is a small-world, not scale-free, network. Proceedings of the Royal Society B: Biological Sciences, 273(1585), 503–511.

45. Just, M. A., Carpenter, P. A., Keller, T. A., Eddy, W. F., & Thulborn, K. R. (1996). Brain activation modulated by sentence comprehension. Science, 274(5284), 114–116.

46. Karbe, H., Thiel, A., Weber-Luxenburger, G., Herholz, K., Kessler, J., & Heiss, W.-D. (1998). Brain plasticity in poststroke aphasia: what is the contribution of the right hemisphere? Brain and language, 64(2), 215–230.

47. Keator, L. M., Yourganov, G., Basilakos, A., Hillis, A. E., Hickok, G., Bonilha, L., … & Fridriksson, J. (2021). Independent contributions of structural and functional connectivity: Evidence from a stroke model. Network Neuroscience, 5(4), 911–928.

48. Keator, L. M., Yourganov, G., Faria, A. V., Hillis, A. E., & Tippett, D. C. (2022). Application of the dual stream model to neurodegenerative disease: evidence from a multivariate classification tool in primary progressive aphasia. Aphasiology, 36(5), 618–647.

49. Kertesz, A. (2007). Western Aphasia Battery--Revised.

50. Kertesz, A. (2022). The Western Aphasia Battery: A systematic review of research and clinical applications. Aphasiology, 36(1), 21–50.

51. Kibby, M. Y., Kroese, J. M., Krebbs, H., Hill, C. E., & Hynd, G. W. (2009). The pars triangularis in dyslexia and ADHD: A comprehensive approach. Brain and language, 111(1), 46–54.

52. Kilpatrick, L. A., Suyenobu, B. Y., Smith, S. R., Bueller, J. A., Goodman, T., Creswell, J. D., … Naliboff, B. D. (2011). Impact of mindfulness-based stress reduction training on intrinsic brain connectivity. Neuroimage, 56(1), 290–298.

53. Kümmerer, D., Hartwigsen, G., Kellmeyer, P., Glauche, V., Mader, I., Klöppel, S., … Saur, D. (2013). Damage to ventral and dorsal language pathways in acute aphasia. Brain, 136(2), 619–629.

54. Labache, L., Joliot, M., Saracco, J., Jobard, G., Hesling, I., Zago, L., … Mazoyer, B. (2019). A SENtence Supramodal Areas AtlaS (SENSAAS) based on multiple task-induced activation mapping and graph analysis of intrinsic connectivity in 144 healthy right-handers. Brain Structure and Function, 224(2), 859–882.

55. LaCroix, A. N., James, E., & Rogalsky, C. (2021). Neural resources supporting language production vs. comprehension in chronic post-stroke aphasia: a meta-analysis using activation likelihood estimates. Frontiers in human neuroscience, 605.

56. Lee, M. H., Smyser, C. D., & Shimony, J. S. (2013). Resting-state fMRI: a review of methods and clinical applications. American journal of neuroradiology, 34(10), 1866–1872.

57. Liao, X., Vasilakos, A. V., & He, Y. (2017). Small-world human brain networks: perspectives and challenges. Neuroscience & Biobehavioral Reviews, 77, 286–300.

58. Liu, J., Wang, Q., Liu, F., Song, H., Liang, X., Lin, Z., … Zheng, G. (2017). Altered functional connectivity in patients with post-stroke memory impairment: A resting fMRI study. Experimental and therapeutic medicine, 14(3), 1919–1928.

59. Liu, Y., Liang, M., Zhou, Y., He, Y., Hao, Y., Song, M., … Jiang, T. (2008). Disrupted small-world networks in schizophrenia. Brain, 131(4), 945–961.

60. Marek, S., Tervo-Clemmens, B., Nielsen, A. N., Wheelock, M. D., Miller, R. L., Laumann, T. O., … & Dosenbach, N. U. (2019). Identifying reproducible individual differences in childhood functional brain networks: An ABCD study. Developmental cognitive neuroscience, 40, 100706.

61. Margulies, D. S., & Petrides, M. (2013). Distinct parietal and temporal connectivity profiles of ventrolateral frontal areas involved in language production. Journal of Neuroscience, 33(42), 16846–16852.

62. Maslov, S., & Sneppen, K. (2002). Specificity and stability in topology of protein networks. Science, 296(5569), 910–913.

63. Matchin, W., den Ouden, D.-B., Hickok, G., Hillis, A. E., Bonilha, L., & Fridriksson, J. (2022). The Wernicke conundrum revisited: evidence from connectome-based lesion-symptom mapping. Brain, 145(11), 3916–3930.

64. Mazrooyisebdani, M., A. Nair, V., Garcia-ramos, C., & Prabhakaran, V. (2018). Abstract TP145: Evolution of Language Network Plasticity Post Stroke Based on Graph Theory. Stroke, 49(Suppl_1), ATP145–ATP145.

65. Mesulam, M.-M., Rogalski, E. J., Wieneke, C., Hurley, R. S., Geula, C., Bigio, E. H., … Weintraub, S. (2014). Primary progressive aphasia and the evolving neurology of the language network. Nature Reviews Neurology, 10(10), 554–569.

66. Montoya, J. M., & Solé, R. V. (2002). Small world patterns in food webs. Journal of theoretical biology, 214(3), 405–412.

67. Muller, A. M., & Meyer, M. (2014). Language in the brain at rest: new insights from resting state data and graph theoretical analysis. Frontiers in human neuroscience, 8, 228.

68. Napadow, V., LaCount, L., Park, K., As-Sanie, S., Clauw, D. J., & Harris, R. E. (2010). Intrinsic brain connectivity in fibromyalgia is associated with chronic pain intensity. Arthritis & Rheumatism, 62(8), 2545–2555.

69. Noble, S., Scheinost, D., Finn, E. S., Shen, X., Papademetris, X., McEwen, S. C., … Cadenhead, K. S. (2017). Multisite reliability of MR-based functional connectivity. Neuroimage, 146, 959–970.

70. Novén, M., Schremm, A., Horne, M., & Roll, M. (2021). Cortical thickness and surface area of left anterior temporal areas affects processing of phonological cues to morphosyntax. Brain Research, 1750, 147150.

71. Pillay, S. B., Binder, J. R., Humphries, C., Gross, W. L., & Book, D. S. (2017). Lesion localization of speech comprehension deficits in chronic aphasia. Neurology, 88(10), 970–975.

72. Postman-Caucheteux, W. A., Birn, R. M., Pursley, R. H., Butman, J. A., Solomon, J. M., Picchioni, D., … Braun, A. R. (2010). Single-trial fMRI shows contralesional activity linked to overt naming errors in chronic aphasic patients. Journal of cognitive neuroscience, 22(6), 1299–1318.

73. Price, C. J. (2012). A review and synthesis of the first 20 years of PET and fMRI studies of heard speech, spoken language and reading. Neuroimage, 62(2), 816–847.

74. Raichle, M. E. (2015). The brain’s default mode network. Annual review of neuroscience, 38, 433–447.

75. Raichle, M. E., MacLeod, A. M., Snyder, A. Z., Powers, W. J., Gusnard, D. A., & Shulman, G. L. (2001). A default mode of brain function. Proceedings of the national academy of Sciences, 98(2), 676–682.

76. Reuter, F., Zaaraoui, W., Crespy, L., Faivre, A., Rico, A., Malikova, I., … & Audoin, B. (2011). Cognitive impairment at the onset of multiple sclerosis: relationship to lesion location. Multiple Sclerosis Journal, 17(6), 755–758.

77. Richter, M., Miltner, W. H., & Straube, T. (2008). Association between therapy outcome and right-hemispheric activation in chronic aphasia. Brain, 131(5), 1391–1401.

78. Rilling, J. K., Glasser, M. F., Preuss, T. M., Ma, X., Zhao, T., Hu, X., & Behrens, T. E. (2008). The evolution of the arcuate fasciculus revealed with comparative DTI. Nature neuroscience, 11(4), 426–428.

79. Richter, M., Miltner, W. H., & Straube, T. (2008). Association between therapy outcome and right-hemispheric activation in chronic aphasia. Brain, 131(5), 1391–1401.

80. Robson, H., Zahn, R., Keidel, J. L., Binney, R. J., Sage, K., & Lambon Ralph, M. A. (2014). The anterior temporal lobes support residual comprehension in Wernicke’s aphasia. Brain, 137(3), 931–943.

81. Rogalsky, C., Basilakos, A., Rorden, C., Pillay, S., LaCroix, A. N., Keator, L., … Fridriksson, J. (2022). The neuroanatomy of speech processing: A large-scale lesion study. Journal of cognitive neuroscience, 34(8), 1355–1375.

82. Rogalsky, C., & Hickok, G. (2009). Selective attention to semantic and syntactic features modulates sentence processing networks in anterior temporal cortex. Cerebral Cortex, 19(4), 786–796.

83. Roger, E., Pichat, C., Torlay, L., David, O., Renard, F., Banjac, S., … Kahane, P. (2019). Hubs disruption in mesial temporal lobe epilepsy. A resting-state fMRI study on a language-and-memory network. Human brain mapping.

84. Ross, L. A., McCoy, D., Wolk, D. A., Coslett, H. B., & Olson, I. R. (2010). Improved proper name recall by electrical stimulation of the anterior temporal lobes. Neuropsychologia, 48(12), 3671–3674.

85. Saur, D., Kreher, B. W., Schnell, S., Kümmerer, D., Kellmeyer, P., Vry, M.-S., … Abel, S. (2008). Ventral and dorsal pathways for language. Proceedings of the national academy of Sciences, 105(46), 18035–18040.

86. Saur, D., Schelter, B., Schnell, S., Kratochvil, D., Küpper, H., Kellmeyer, P., … Lange, R. (2010). Combining functional and anatomical connectivity reveals brain networks for auditory language comprehension. Neuroimage, 49(4), 3187–3197.

87. Schöpf, V., Kasess, C., Lanzenberger, R., Fischmeister, F., Windischberger, C., & Moser, E. (2010). Fully exploratory network ICA (FENICA) on resting-state fMRI data. Journal of neuroscience methods, 192(2), 207–213.

88. Shulman, G. L., Fiez, J. A., Corbetta, M., Buckner, R. L., Miezin, F. M., Raichle, M. E., & Petersen, S. E. (1997). Common blood flow changes across visual tasks: II. Decreases in cerebral cortex. Journal of cognitive neuroscience, 9(5), 648–663.

89. Spitsyna, G., Warren, J. E., Scott, S. K., Turkheimer, F. E., & Wise, R. J. (2006). Converging language streams in the human temporal lobe. Journal of Neuroscience, 26(28), 7328–7336.

90. Stam, C., De Haan, W., Daffertshofer, A., Jones, B., Manshanden, I., van Cappellen van Walsum, A.-M., … Van Dijk, B. (2009). Graph theoretical analysis of magnetoencephalographic functional connectivity in Alzheimer’s disease. Brain, 132(1), 213–224.

91. Tedeschi, G., Russo, A., Conte, F., Corbo, D., Caiazzo, G., Giordano, A., … Tessitore, A. (2016). Increased interictal visual network connectivity in patients with migraine with aura. Cephalalgia, 36(2), 139–147.

92. Tessitore, A., De Micco, R., Giordano, A., di Nardo, F., Caiazzo, G., Siciliano, M., … Tedeschi, G. (2017). Intrinsic brain connectivity predicts impulse control disorders in patients with Parkinson’s disease. Movement Disorders, 32(12), 1710–1719.

93. Thye, M., & Mirman, D. (2018). Relative contributions of lesion location and lesion size to predictions of varied language deficits in post-stroke aphasia. NeuroImage: Clinical, 20, 1129–1138.

94. Tie, Y., Rigolo, L., Norton, I. H., Huang, R. Y., Wu, W., Orringer, D., … Golby, A. J. (2014). Defining language networks from resting-state fMRI for surgical planning—a feasibility study. Human brain mapping, 35(3), 1018–1030.

95. Tombaugh, T. N., & McIntyre, N. J. (1992). The mini-mental state examination: a comprehensive review. Journal of the American Geriatrics Society, 40(9), 922–935.

96. Trimmel, K., van Graan, A. L., Caciagli, L., Haag, A., Koepp, M. J., Thompson, P. J., & Duncan, J. S. (2018). Left temporal lobe language network connectivity in temporal lobe epilepsy. Brain, 141(8), 2406–2418.

97. Van Den Heuvel, M. P., & Pol, H. E. H. (2010). Exploring the brain network: a review on resting-state fMRI functional connectivity. European neuropsychopharmacology, 20(8), 519–534.

98. Van Ettinger-Veenstra, H. M., Ragnehed, M., Hällgren, M., Karlsson, T., Landtblom, A. M., Lundberg, P., & Engström, M. (2010). Right-hemispheric brain activation correlates to language performance. Neuroimage, 49(4), 3481–3488.

99. Vandenberghe, R., Nobre, A. C., & Price, C. (2002). The response of left temporal cortex to sentences. Journal of cognitive neuroscience, 14(4), 550–560.

100. Vigneau, M., Beaucousin, V., Hervé, P.-Y., Duffau, H., Crivello, F., Houde, O., … Tzourio-Mazoyer, N. (2006). Meta-analyzing left hemisphere language areas: phonology, semantics, and sentence processing. Neuroimage, 30(4), 1414–1432.

101. Vincent, J. L., Kahn, I., Snyder, A. Z., Raichle, M. E., & Buckner, R. L. (2008). Evidence for a frontoparietal control system revealed by intrinsic functional connectivity. Journal of neurophysiology, 100(6), 3328–3342.

102. Wallentin, M., Nielsen, A. H., Vuust, P., Dohn, A., Roepstorff, A., & Lund, T. E. (2011). BOLD response to motion verbs in left posterior middle temporal gyrus during story comprehension. Brain and language, 119(3), 221–225.

103. Walsh, M. J., Baxter, L. C., Smith, C. J., & Braden, B. B. (2019). Age group differences in executive network functional connectivity and relationships with social behavior in men with autism spectrum disorder. Research in autism spectrum disorders, 63, 63–77.

104. Watts, D. J., & Strogatz, S. H. (1998). Collective dynamics of ‘small-world’networks. Nature, 393(6684), 440–442.

105. Wise, R. J., Scott, S. K., Blank, S. C., Mummery, C. J., Murphy, K., & Warburton, E. A. (2001). Separate neural subsystems withinWernicke’s area’. Brain, 124(1), 83–95.

106. Wolmetz, M., Poeppel, D., & Rapp, B. (2011). What does the right hemisphere know about phoneme categories? Journal of cognitive neuroscience, 23(3), 552–569.

107. Xiao, Y., Friederici, A. D., Margulies, D. S., & Brauer, J. (2016). Longitudinal changes in resting-state fMRI from age 5 to age 6 years covary with language development. Neuroimage, 128, 116–124.

108. Xing, S., Lacey, E. H., Skipper-Kallal, L. M., Jiang, X., Harris-Love, M. L., Zeng, J., & Turkeltaub, P. E. (2016). Right hemisphere grey matter structure and language outcomes in chronic left hemisphere stroke. Brain, 139(1), 227–241.

109. Xu, L., Huang, L., Cui, W., & Yu, Q. (2020). Reorganized functional connectivity of language centers as a possible compensatory mechanism for basal ganglia aphasia. Brain Injury, 34(3), 430–437.

110. Yang, M., Li, Y., Li, J., Yao, D., Liao, W., & Chen, H. (2017). Beyond the arcuate fasciculus: damage to ventral and dorsal language pathways in aphasia. Brain topography, 30(2), 249–256.

111. Zhang, J., Kucyi, A., Raya, J., Nielsen, A. N., Nomi, J. S., Damoiseaux, J. S., … Whitfield-Gabrieli, S. (2021). What have we really learned from functional connectivity in clinical populations? Neuroimage, 242, 118466.

112. Zhao, Y., Ralph, M. A. L., & Halai, A. D. (2018). Relating resting-state hemodynamic changes to the variable language profiles in post-stroke aphasia. NeuroImage: Clinical, 20, 611–619.

113. Zhu, D., Chang, J., Freeman, S., Tan, Z., Xiao, J., Gao, Y., & Kong, J. (2014). Changes of functional connectivity in the left frontoparietal network following aphasic stroke. Frontiers in Behavioral Neuroscience, 8, 167.

114. Zhu, H., Zhou, P., Alcauter, S., Chen, Y., Cao, H., Tian, M., … Zhao, X. (2016). Changes of intranetwork and internetwork functional connectivity in Alzheimer’s disease and mild cognitive impairment. Journal of neural engineering, 13(4), 046008.

